# Capturing single-copy nuclear genes, organellar genomes, and nuclear ribosomal DNA from deep genome skimming data for plant phylogenetics: A case study in Vitaceae

**DOI:** 10.1101/2021.02.25.432805

**Authors:** Bin-Bin Liu, Zhi-Yao Ma, Chen Ren, Richard G.J. Hodel, Miao Sun, Xiu-Qun Liu, Guang-Ning Liu, De-Yuan Hong, Elizabeth A. Zimmer, Jun Wen

## Abstract

With the decreasing cost and availability of many newly developed bioinformatics pipelines, next-generation sequencing (NGS) has revolutionized plant systematics in recent years. Genome skimming has been widely used to obtain high-copy fractions of the genomes, including plastomes, mitochondrial DNA (mtDNA), and nuclear ribosomal DNA (nrDNA). In this study, through simulations, we evaluated optimal (minimum) sequencing depth and performance for recovering single-copy nuclear genes (SCNs) from genome skimming data, by subsampling genome resequencing data and generating 10 datasets with different sequencing coverage *in silico*. We tested the performance of the four datasets (plastome, nrDNA, mtDNA, and SCNs) obtained from genome skimming based on phylogenetic analyses of the *Vitis* clade at the genus-level and Vitaceae at the family-level, respectively. Our results showed that optimal minimum sequencing depth for high-quality SCNs assembly via genome skimming was about 10× coverage. Without the steps of synthesizing baits and enrichment experiments, we showcase that deep genome skimming (DGS) is effective for capturing large datasets of SCNs, in addition to plastomes, mtDNA, and entire nrDNA repeats, and may serve as an economical alternative to the widely used target enrichment Hyb-Seq approach.

## 1 Introduction

Genome skimming has often been used to target the high-copy fractions of genomes including plastomes, mitochondrial genomes (mitogenomes), and nuclear ribosomal DNA (nrDNA) repeats (Straub et al., 2012; Dodsworth, 2015; Zhang et al., 2015; Thode et al., 2020), and these datasets have been widely used for inferring phylogenies in many recent studies. For example, the chloroplast genome has been widely utilized for inferring the phylogenetic relationships at various levels (Bock et al., 2014; Zhang et al., 2015; Valcárcel & Wen, 2019; Zhang et al., 2019; Wang et al., 2020), clarifying generic and species delimitations (Wen et al., 2018a; Liu et al., 2019; 2020a; 2020b), as well as acting as an ultra-barcode in plants (Kane et al., 2012; Hollingsworth et al., 2016). The uniparental (mostly maternal, rarely paternal) inheritance and non-recombinant nature of the plastomes make them the ideal marker for tracking the maternal (rarely paternal) history, providing useful evidence to untangle hybridization events in plants (Rieseberg & Soltis, 1991; Sun et al., 2015; Folk et al., 2017; Vargas et al., 2017; Morales-Briones et al., 2018). The mitogenome has not been widely used as a source of phylogenetic data in plants due to its low nucleotide substitution rates (Palmer & Herbon, 1988; Palmer, 1990), concerns over the impact of RNA editing on phylogenetic reconstruction (Sloan et al., 2009; Mower et al., 2012; Wu et al., 2021), and the supposedly shared evolutionary history with plastomes (Rieseberg & Soltis, 1991; Olson & McCauley, 2000). The mitogenome nevertheless has been useful in phylogenetic estimation at higher taxonomic levels, e.g., at the family level in Rubiaceae (Rydin et al., 2017) and Vitaceae (Zhang et al., 2015). Some regions of the nrDNA repeat, especially the internal transcribed spacer (ITS) and sometimes also the external transcribed spacer (ETS) have been widely used for lower-level phylogenetic reconstruction in flowering plants (Baldwin et al., 1995; Álvarez & Wendel, 2003; Soltis et al., 2008). Recently, the entire nrDNA repeats including ETS, 18S, ITS1, 5.8S, ITS2, and 26S regions have been assembled from genome skimming data, and depending on the region of the repeat that has been utilized, has also been effective in providing phylogenetic resolution at shallow evolutionary levels (for example, in the Rosaceae: Liu et al., 2019, 2020a, 2020b). Hence, genome skimming has been a valuable approach for providing genomic data for phylogenetic inferences.

Because genome skimming data are generated from the total genomic DNA, the organellar genomes (plastome and mitogenome) only account for a small portion of the reads, e.g., only 4-5% of the data accounting for plastomes (Straub et al., 2012), indicating the underutilization of the genome skimming data, which may have potential for the discovery of nuclear markers. Several recent studies have demonstrated the promise of genome skimming in exploring single-copy nuclear genes (SCNs) (Berger et al., 2017; Vargas et al., 2019). Berger et al. (2017) obtained three low-copy nuclear *CYC*-like genes for detailed evo-devo analysis in a 2× to 3.5× coverage genome skimming dataset. Moreover, Vargas et al. (2019) designed 354 nuclear loci with the combination of *MarkerMiner* (Chamala et al., 2015) and their custom-designed tool *GoldFinder* using five transcriptomes of Lecythidoideae. All these 354 loci were captured *in silico* from a prior genome skimming data set, opening a new window for using genome skimming data to screen nuclear loci. However, Vargas et al. (2019) used only the reference-guided assembly method for targeting the nuclear genes from low nuclear genomic coverage, making it unsuitable for assessing orthology. These two case studies showed the potential of genome skimming data in recovering SCNs.

Harboring genetic information from both parents, single/low-copy nuclear genes have been utilized as valuable markers for phylogenetic inferences in angiosperms (Zhang et al., 2012; Zimmer & Wen, 2012, 2015). Next-generation sequencing (NGS) has provided an opportunity for capturing a large number of nuclear genes, addressing problems unresolvable using traditional molecular systematics approaches (e.g., Léveillé-Bourret et al., 2018; Herrando-Moraira et al., 2019). Large datasets of nuclear genes have facilitated the use of species tree methods based on multispecies coalescent models (Mirarab & Warnow, 2015; Edwards et al., 2016), which have greatly increased the accuracy of phylogenetic inference (McCormack et al., 2009; Smith et al., 2015). Among the genome-scale methods developed to date, target enrichment, also known as Hyb-Seq, has been shown as the most efficient and cost-effective approach for obtaining large datasets of single-copy nuclear genes (SCNs) for plant systematics (Lemmon et al., 2012; Mandel et al., 2014; Weitemier et al., 2014; Dodsworth et al., 2019). Targeted nuclear sequences from Hyb-Seq have been corroborated to be effective for providing greater phylogenetic resolution both at shallow and deep levels (Villaverde et al., 2018; Kleinkopf et al., 2019; Li et al., 2019; Ma et al., 2021). Due to its good performance with degraded DNA from silica gel-dried and herbarium specimens (Weitemier et al., 2014; Villaverde et al., 2018; Wang et al., 2021), Hyb-Seq has gained popularity in recent phylogenomic studies, unlike whole-genome sequencing (WGS) and transcriptome sequencing (RNAseq) that require fresh or flash-frozen materials (Xiang et al., 2017). However, because the 80-120 bp RNA baits are required for hybridizing experiments to library inserts in target enrichment methods, a balance is needed between selecting genomic regions variable enough to infer phylogenies and those conserved enough to ensure sequence recovery; and such balance has greatly limited the number of SCNs designed from closely-related genomes and/or transcriptomes. In addition, the high costs of generating customized baits and the complex experimental procedures have also impeded the utilization of this method in many labs. In particular, it is practically difficult in many developing countries, without easy access to synthesized baits.

Given the promise of genome skimming for recovering SCNs, we used simulations to subsample genome resequencing data from our previous study (Ma et al., 2018) to explore the optimal sequencing depths for obtaining sufficient SCNs for plant phylogenetics. We designed two study cases in the grape family: (1) a family-level case in Vitaceae, a medium-sized plant family with about 950 species belonging to 16 extant genera that include dominant climbers in both tropical and temperate zones (Wen et al., 2007; 2018b), and (2) a genus-level case in the grapevine genus *Vitis* L., consisting of c. 70 species predominantly from the Northern Hemisphere (Liu et al., 2016; Wen et al., 2018a). Ma et al. (2018) used SNP calling of genome resequencing data for 41 samples of *Vitis*. As the dataset has high 20× coverage on average, it provides sufficient raw data to explore four genomic counterparts: plastomes, mitogenomes, nrDNA, and SCNs for phylogenetic reconstruction. Furthermore, its 20× coverage represents a good opportunity to randomly generate 10 subsamples with different sequencing depths for testing the optimal sequencing depths to capture sufficient SCNs for phylogenetic analyses. We also tested the utility for SCN generation of one low-coverage genome skimming data set that originally had been sequenced by Zhang et al. (2015) to recover organelle DNA data in Vitaceae.

## 2 Materials & Methods

### 2.1 Sequencing depths for each case

For the genus-level case, we used the genome resequencing dataset of *Vitis* by Ma et al. (2018), representing 41 species sequenced using Illumina Hi-Seq (NCBI Short Read Archive SRP161488 under the BioProject PRJNA490319). Detailed species and voucher information can be found in Ma et al. (2018) and Table S1. The sequencing depths of these 41 datasets ranged from 17.5× to 33.4× coverage with approximately 20× coverage on average (Table S1), assuming an estimated genome size of around 487 Mb based on the *Vitis vinifera* L. genome (Jaillon et al., 2007).

For the family-level case, we used the low-coverage genome skimming dataset of 27 Vitaceae species generated on an Illumina Next-Seq instrument by Zhang et al. (2015) to test the effectiveness of simulation results in Vitaceae. All raw data were downloaded from the GenBank with the BioProject accession number PRJNA298058 and the sequencing depths ranged from 4× to 7.4× coverage (average 5.6× coverage). Detailed species and voucher information are available in Zhang et al. (2015) and Table S2.

### 2.2 Capturing single-copy nuclear genes *in silico* via genome skimming

#### 2.2.1 Data subset creation

For the dataset of 41 genome resequencing samples of *Vitis*, we used the python script *randomReadSubSample.py* (Piro et al., 2017) to make random draws from the raw data files. Nine different subset sequencing depths were generated, 2× (10%), 4× (20%), 6× (30%), 8× (40%), 10× (50%), 12× (60%), 14× (70%), 16× (80%), and 18× (90%), because we expected the minimum sequencing depth for success would fall within this range.

#### 2.2.2 Single-copy nuclear marker development

Targeted nuclear genes were selected from the coding regions of *Vitis vinifera* (GenBank assembly accession: GCA_000003745.2). The coding sequences were first submitted to MarkerMiner v.1.0 (Chamala et al., 2015) to identify the putative single-copy genes. The genome of *V. vinifera* as a proteome reference has been integrated into MarkerMiner, and the default settings of the program were followed, except that the minimum sequence length was set as “600 bp” in order to acquire more candidate genes. The resulting genes were then filtered by successively BLASTing (Altschul et al., 1990, 1997; Camacho et al., 2009) them against four available *Vitis* genomes (*V. aestivalis* Michx., GCA_001562795.1; *V. cinirea* (Engelm.) Millardet × *V. riparia* Michx., GCA_001282645.1; *V. riparia*, GCA_004353265.1; and *V. vinifera*, GCA_000003745.2) in Geneious Prime (Kearse et al., 2012), with the parameters settings in the Megablast program (Morgulis et al., 2008) as a maximum of 60 hits, a maximum E-value of 1 × 10^−10^, a linear gap cost, a word size of 28, and scores of 1 for match and −2 for mismatch in alignments. We first excluded the genes with mean coverage > 1.1 for alignments, which generally would suggest potential paralogy of the genes and/or the presence of highly repeated elements in the sequences. The remaining alignments were further visually examined to exclude those genes receiving multiple hits with long overlapping but different sequences during BLASTing. It should be noted that the alignments with mean coverage between 1.0 and 1.1 were generally caused by the presence of tiny pieces of flanking intron sequences in the alignments. These fragments were still accepted as an SCN here. After the filtration, the remaining genes were used as references in the following gene assembly.

#### 2.2.3 Targeting nuclear single-copy genes

We used Trimmomatic v. 0.39 (Bolger et al., 2014) for quality trimming and adapter clipping, with removing the leading/trailing low quality or below quality three bases, scanning the read with a 4-base wide sliding window, cutting when the average quality per base drops below 14, and dropping reads below 36 bases long. Subsequently, the results were quality-checked using FastQC v. 0.11.9 (Andrews, 2018). The HybPiper pipeline v. 1.3.1 (Johnson et al., 2016) was used for targeting SCNs with default settings; BWA v. 0.7.1 (Li & Durbin, 2009) to align and distribute reads to target genes; SPAdes v. 3.15.0 (Bankevich et al., 2012) with a coverage cutoff value of 5 to assemble reads to contigs; and Exonerate v. 2.2.0 (Slater & Birney, 2005) to align assembled contigs to target sequences and determine exon-intron boundaries. Python and R scripts included in the HybPiper pipeline (Johnson et al., 2016) were used to retrieve the recovered gene sequences, and to summarize and visualize the recovery efficiency. The final alignment of the SCNs from the 10 subsampling datasets are available from the Dryad Digital Repository: https://doi.org/10.5061/dryad.b2rbnzsd7 (Liu et al., 2021).

### 2.3 Assembly of chloroplast genome and nrDNA repeats by a successive method

To obtain high-quality chloroplast genomes and nrDNA repeats, a two-step strategy was used for assembly. NOVOPlasty v. 4.3.1 (Dierckxsens et al., 2016) was applied first to assemble the plastomes with high-quality raw data and then we used the successive assembly approach by Zhang et al. (2015), combining the reference-based and the *de novo* assembly methods to assemble the remaining low-quality samples. With the *de novo* assembly and a seed-and-extend algorithm, NOVOPlasty was the least laborious approach and resulted in the most accurate plastomes; however, this program needs sufficient high-quality raw reads without gaps to cover the whole plastome. The whole plastomes assembled from NOVOPlasty then could be used as references for assembling the remaining samples. The successive method provided us with a good approach to obtain relatively accurate and nearly complete plastomes with or without gaps from lower-coverage raw data. Due to the sensitivity of Bowtie2 v. 2.4.2 (Langmead & Salzberg, 2012) to the reference, this successive method needs a closely related reference sequence with more time and RAM requirement. Because the nuclear ribosomal DNA copies are arranged in tandem repeats, we tentatively treated the complete nrDNA as circular in order to use the chloroplast genome assembly software (NOVOPlasty) for the rDNA assembly, and the steps we used were nearly the same as the assembly procedure of plastomes as described above. The detailed procedure has been described in several recent studies (Zhang et al., 2015; Liu et al., 2019, 2020a, 2020b). All the assembled plastomes and nrDNA repeats have been submitted to GenBank with the accession numbers listed in Table S1 & S2.

### 2.4 Assembly of mitogenome genes

Given the highly variable structure and the recurrent rearrangements in plant mitochondrial genomes, we extracted the genic portion of *Vitis vinifera* mitogenome for phylogenetic analyses by Geneious Prime (Kearse et al., 2012), in which 37 mitochondrial origin protein-coding genes (38 genes, as *rps19* has two functional full gene copies) were included (Goremykin et al., 2009). It should be noted that the mitochondrial origin rRNA, tRNA, and hypothetical genes (Goremykin et al., 2009) were not included in the targeted gene list. All these 38 genes in *Vitis vinifera* were used as the reference in HybPiper (Johnson et al., 2016) to capture the mitogenes for the other species with the coverage cutoff 5 and the other parameters by default. The final alignments are available from the Dryad Digital Repository (Data S12&S13): https://doi.org/10.5061/dryad.b2rbnzsd7 (Liu et al., 2021).

### 2.5 Sequence annotation and alignment

The assembled plastid genomes from the low-coverage and high-coverage datasets were annotated using PGA (Qu et al., 2019) with a closely related plastome (MT267294) downloaded from GenBank as the reference, and the results of automated annotation checked manually. The coding sequences of plastomes were translated into proteins to check the start and stop codons manually in Geneious Prime (Kearse et al., 2012). The custom annotations in the GenBank format were converted into the FASTA and five-column feature tables file required by NCBI submission using GB2sequin (Lehwark & Greiner, 2019).

The entire nrDNA sequence includes six regions, ETS, 18S, ITS1, 5.8S, ITS2, and 26S, it should be noted that the nontranscribed spacer region between the 26S and the ETS was excluded here due to the ambiguous alignment. All sequences from plastome, mitogenome, nrDNA repeats, and SCNs assembled here were aligned separately by MAFFT v. 7.475 (Nakamura et al., 2018) with default parameters. Specifically, as the sequences of the two IR regions of the plastome in each assession of the grape family were completely or nearly identical, only one copy of the inverted repeat (IR) region was included for the whole plastome phylogenetic analyses. To reduce the systematic errors produced by poor alignment, we used trimAL v. 1.2 (Capella-Gutiérrez et al., 2009) to trim the alignment of these sequences, in which all columns with gaps in more than 20% of the sequences or with a similarity score lower than 0.001 were removed. Each aligned sequence of SCNs, plastid coding sequences (CDS), nrDNA regions, and mtDNA genes was concatenated by AMAS v. 1.0 (Borowiec, 2016), respectively, and the resulting alignment summaries of each dataset have been used to estimate the partition of each gene.

### 2.6 Phylogenetic analyses

Bayesian inferences (BI) were run for the whole plastome of *Vitis*, plastid CDS of Vitaceae, mtDNA, and nrDNA dataset at the genus level of *Vitis* and the family level of Vitaceae separately. The best-fit partitioning schemes and/or nucleotide substitution models for each dataset were estimated using PartitionFinder2 (Stamatakis, 2006; Lanfear et al., 2016), under the corrected Akaike information criterion (AICc) and linked branch lengths, as well as with greedy (Lanfear et al., 2012) for the nrDNA dataset and rcluster (Lanfear et al., 2014) algorithm options for plastid CDS and mtDNA dataset. The partitioning schemes and evolutionary model for each subset were used for the downstream Bayesian Inference (BI) analyses. The BI tree was performed with MrBayes 3.2.7 (Ronquist et al., 2012). The Markov chain Monte Carlo (MCMC) analyses were run for 100,000,000 generations. Trees were sampled at every 2,000 generations with the first 25% discarded as burn-in. The remaining trees were used to build a 50% majority-rule consensus tree. The stationarity was regarded to be reached when the average standard deviation of split frequencies remained below 0.01. The BI tree was visualized using Geneious Prime (Kearse et al., 2012).

For the SCNs, we inferred phylogenies using both concatenation and species tree methods. For the concatenation analysis, we used the aforementioned PartitionFinder2 (Stamatakis, 2006; Lanfear et al., 2016) to estimate the best partitioning schemes for each gene with the parameters same as above except for using the rcluster algorithm option, and the resulted schemes were then used to infer Maximum Likelihood (ML) trees with RAxML 8.2.12 (Stamatakis, 2014). For estimating the coalescent species tree, we searched for the best-scoring ML trees and performed 100 rapid bootstraps employing the option “-f a” in RAxML 8.2.12 (Stamatakis, 2014), using an independent GTRGAMMA model for each of the 887 SCNs. The gene trees were then used to infer a coalescent-based species tree with ASTRAL-III (Zhang et al., 2018), which infers a species tree from gene trees accounting for the incongruence produced by incomplete lineage sorting (ILS). Each of the gene trees was rooted and low support branches (≤ 10) were contracted by Newick Utilities (Junier & Zdobnov, 2010), since collapsing gene tree nodes with BS support less than a certain value will help to improve accuracy (Zhang et al., 2018).

Because the phylogeny inferred in this study using 887 SCNs (see the results below) showed some incongruence with that in Ma et al. (2018) based on single nucleotide polymorphisms (SNPs), we employed *phyparts* v. 0.0.1 (Smith et al., 2015) to calculate the amount of conflict among the 887 SCN gene trees by comparing the nuclear gene trees against the ASTRAL species tree. We performed phylogenetic conflict analysis using *phyparts* with a bootstrap support (BS) threshold of 30 (i.e., gene-tree branches/nodes with less than 30% BS were considered uninformative), although BS with 70% in some studies has been used as the cutoff (Stull et al., 2020). The baseline for strong support has been rightfully challenged (Soltis & Soltis, 2003). Nevertheless, it is useful, albeit somewhat arbitrary, for filtering out poorly supported branches, thus alleviating noise in the results of the conflict analysis (Smith et al., 2015). *Phyparts* results were visualized with *phypartspiecharts.py* (by Matt Johnson, available from https://github.com/mossmatters/MJPythonNotebooks/blob/master/phypartspiecharts.py).

## 3 Results

### 3.1 Capturing single-copy nuclear loci

We created ten datasets *in silico* with different coverage levels for the downstream analyses (Table 1). The number of genes recovered increased with the increase of coverage in each dataset (Fig. 1A-J), and all the assembled SCNs have been deposited in Dryad Digital Repository (Data S1-S10) https://doi.org/10.5061/dryad.b2rbnzsd7 (Liu et al., 2021). We obtained 884 SCNs included in more than 95% samples and the 887 genes included in more than 80% samples from the 20× coverage genome resequencing data (Table 1). Balancing missing data and the number of genes, we selected genes with more than 50% samples (≥21 samples) retained for the following phylogenetic analyses. Our result showed that only 31 nuclear genes with 50% samples have been recovered from the 6× coverage data, two from the 4× coverage data, and 0 from the 2× coverage data (Table 1). 618 SCNs have been recovered from the 8× coverage data, 876 SCNs were successfully assembled from the 10× coverage data, 885 SCNs from the 12× coverage data, and all 887 genes with more than 50% samples were recovered from the 14×, 16×, 18×, and 20× coverage (Table 1).

**Fig. 1.**
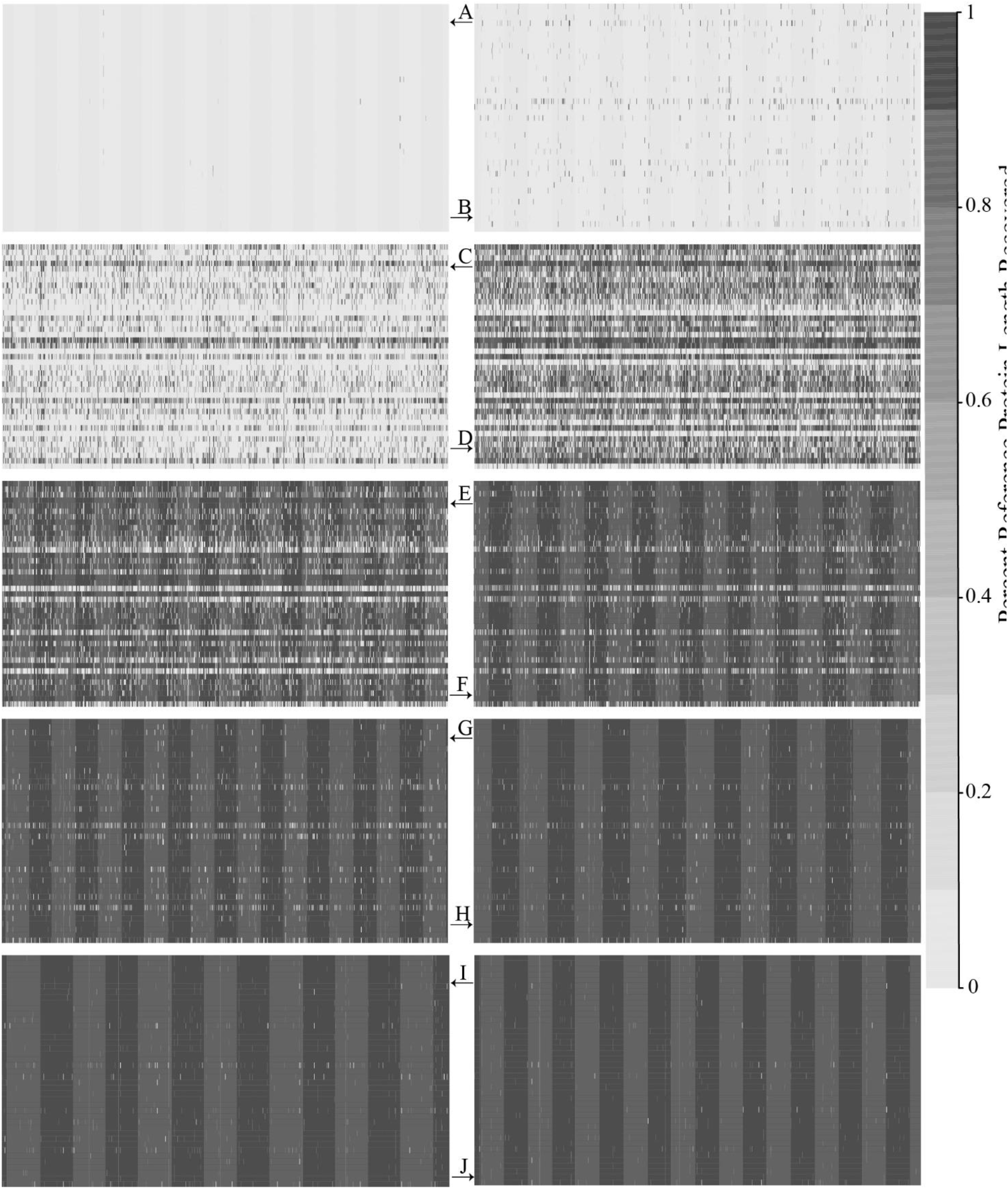
Heat map showing recovery efficiency for 887 genes enriched in *Vitis* recovered by HybPiper. Each column is a gene, and each row is one sample. A, 2× coverage subsampling data; B, 4× coverage subsampling data; C, 6× coverage subsampling data; D, 8× coverage subsampling data; E, 10× coverage subsampling data; F, 12× coverage subsampling data; G, 14× coverage subsampling data; H, 16× coverage subsampling data; I, 18× coverage subsampling data; J, 20× coverage raw data. The shade of gray in the cell is determined by the length of sequence recovered by the pipeline, divided by the length of the reference gene (maximum of 1.0). Full data for each subsample can be found in Dryad Digital Repository Data S1-S10.

**Table 1.** Assembly table for 41 samples in *Vitis* at different sequencing depths

| Sequencing | 95% | 90% | 80% | 70% | 60% | **50%** | 30% | 10% |
| --- | --- | --- | --- | --- | --- | --- | --- | --- |
| Coverage\Percentage | ($\geq 39$)<sup>2</sup> | ($\geq 37$) | ($\geq 33$) | ($\geq 29$) | ($\geq 25$) | ($\geq 21$) | ($\geq 13$) | ($\geq 5$) |
| Recovered |  |  |  |  |  |  |  |  |
| 10% (2X)<sup>1</sup> | 0 | 0 | 0 | 0 | 0 | **0** | 0 | 2 |
| 20% (4X) | 0 | 0 | 0 | 0 | 1 | **2** | 8 | 27 |
| 30% (6X) | 0 | 1 | 3 | 6 | 17 | **31** | 200 | 737 |
| 40% (8X) | 11<sup>3</sup> | 29 | 107 | 249 | 443 | **618** | 849 | 885 |
| **<u>50% (10X)</u>** | **<u>174</u>** | **<u>350</u>** | **<u>632</u>** | **<u>779</u>** | **<u>852</u>** | **<u>876</u>** | **<u>885</u>** | **<u>887</u>** |
| 60% (12X) | 583 | 739 | 853 | 880 | 881 | **885** | 886 | 887 |
| 70% (14X) | 811 | 862 | 879 | 885 | 886 | **887** | 887 | 887 |
| 80% (16X) | 869 | 881 | 885 | 887 | 887 | **887** | 887 | 887 |
| 90% (18X) | 879 | 886 | 887 | 887 | 887 | **887** | 887 | 887 |
| 100% (20X) | 884 | 886 | 887 | 887 | 887 | **887** | 887 | 887 |

Notes: 1. The rows indicate the coverage of genome skimming data *in silico* with the sequencing depths in parenthesis;
2. The columns indicate the number of genes recovered for the related percentage (number) of samples;
3. Eleven nuclear genes with equal to and more than 39 samples were recovered in the 8× genome skimming data *in silico*

The ASTRAL species trees estimated from the different datasets resulted in increased support from these ten datasets *in silico* (Figs, S1-S7). The species tree based on the 20× coverage dataset resolved the phylogenetic relationships among taxa (Fig. 2A), while the species trees from the 2×, 4×, 6×, and 8× coverage data provided limited inference of specific relationships (Figs. S1, S2). However, our results showed that the species tree estimated from the 10× coverage data resulted in a tree as highly supported as that from 20× coverage data, with some differences between these two topologies due to lower support in some clades (Fig. S3). The other four ASTRAL species trees (Figs. S4-S7) presented a highly supported topology, all of which were based on datasets from more than 800 nuclear genes. Our results indicated that 10× coverage genome skimming data was the minimum sequencing depth for recovering sufficient SCNs for the phylogenetic analyses.

**Fig. 2.**
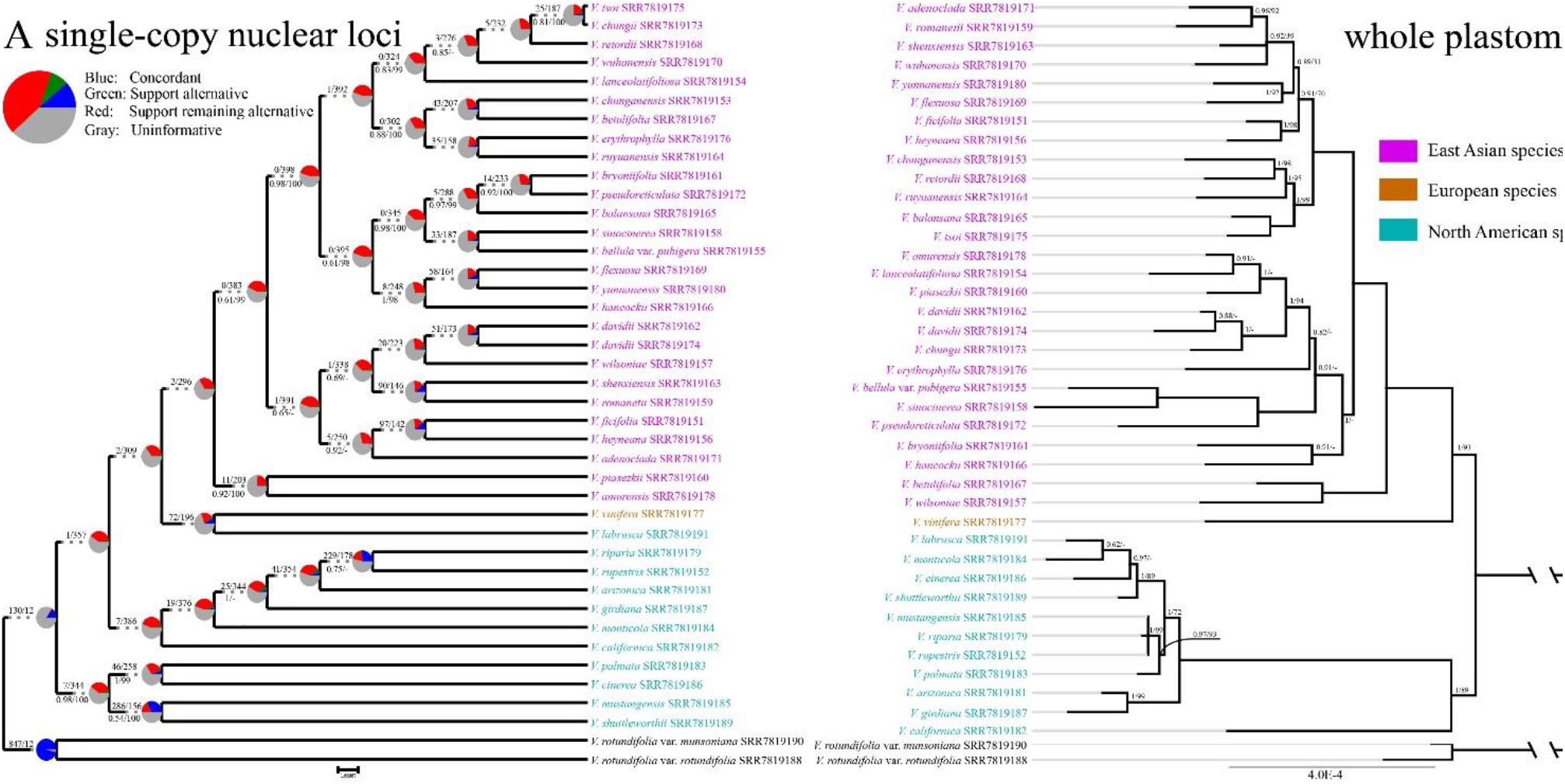
Comparisons of the ASTRAL species tree inferred from 887 SCNs and bayesian trees estimated from the whole plastome of the *Vitis* data. (A) also shows the concordant and conflict of the reduced 523 SCNs., and Pie charts indicate the proportion of genes that agree (blue), support a main alternative topology (green), support the remaining alternatives (red), and are uninformative (gray) for a given node on the underlying topology. Numbers above the nodes show the number of concordant genes/that of conflicting genes (support main alternative + support remaining alternatives). While the number under the nodes indicate the branch support values measuring the support for a quadripartition/maximum likelihood bootstrap support, and all nodes have quadripartition branch support of 1 and bootstrap support of 100 unless noted otherwise. Lines between taxa indicate a conflicting position between these two topologies.

The ASTRAL species tree based on 887 SCNs from the 20× coverage revealed the paraphyly of the North American species in *Vitis* (Fig. 2A), contrasting the results in Ma et al. (2018), which showed the monophyly of this group. Additionally, the analysis of phylogenetic conflict with *phyparts* showed that 130 out of 142 informative SCNs (91.5%) supported this paraphyly of the North American grape species (Fig. 2A). The 4× - 7.4× coverage genome skimming data simulated in our *in silico* analysis resulted in 2-31 SCNs with more than 50% samples, and our empirical analysis of the Vitaceae data from Zhang et al. (2015) did not recover any SCNs with more than 50% samples using a coverage cutoff of 5.

### 3.2 High-copy fractions of genomes: whole plastome, nrDNA, and mtDNA sequences

We used the optimal 10× coverage data aimed for recovering SCNs proposed above to assemble the high-copy fractions of genomes of Vitaceae: whole plastome, nrDNA, and mtDNA sequences, all of which have been well-assembled (Table S1 & Dryad Digital Repository Data S13: https://doi.org/10.5061/dryad.b2rbnzsd7, Liu et al., 2021). Given the low sequence divergence and good alignment of intergenic regions among plastomes in *Vitis*, we used the whole plastome to estimate the phylogeny of *Vitis* (Fig. 2B). The plastid tree resulted in three strongly supported clades, the European clade, the East Asian clade, and the North American clade (Fig. 2B). These three clades were also supported by our mtDNA tree, contrasting with the paraphyly of the North American clade supported by the 887 SCNs (Fig. 2A). The cytonuclear discordance was also detected in the phylogenetic position of the North American species *V. labrusca* L., and this species grouped with other North American species in the plastid and mtDNA trees (Figs. 2B, 4A), while was sister to the European species *V. vinifera* in the nuclear tree (Fig. 2A). Unfortunately, the nrDNA sequences from the *Vitis* data did not provide sufficient informative sites to clarify phylogenetic relationships within *Vitis* (Fig. 3A).

**Fig. 3.**
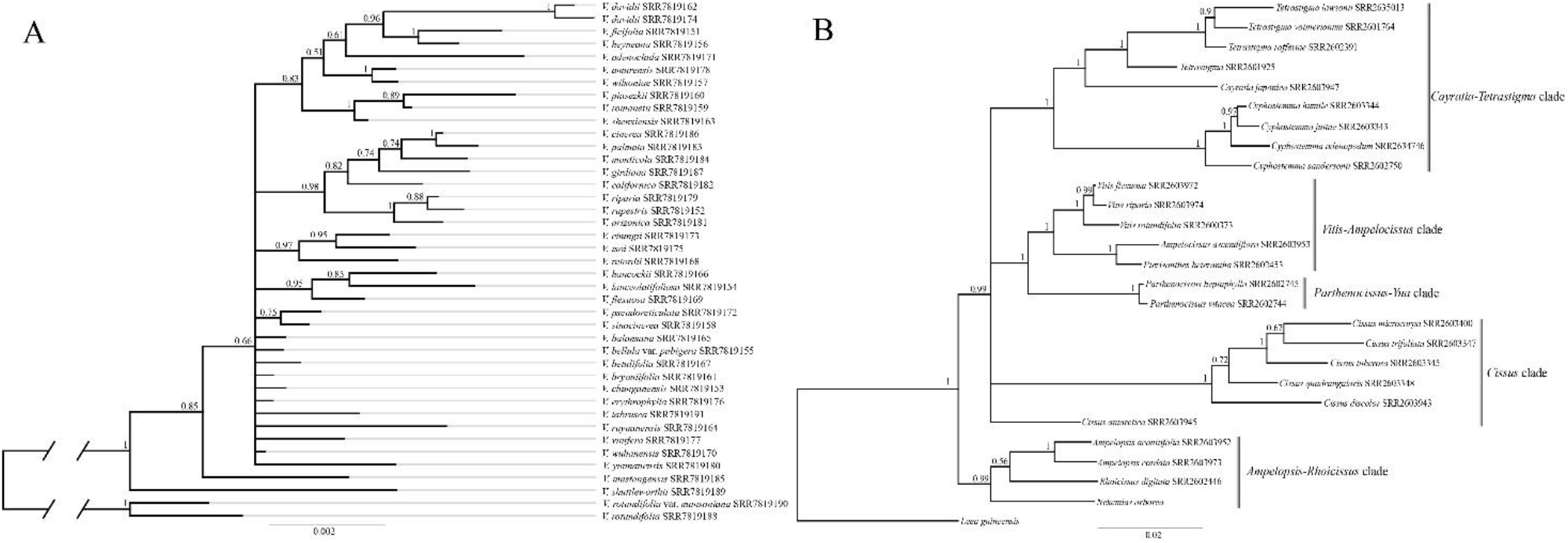
Bayesian trees inferred from nrDNA of *Vitis* (A) and Vitaceae (B) data. The number above the nodes indicate the branch support values measuring the support for the BI posterior probabilities (PP). Scale bars indicate substitutions per site. nrDNA, nuclear ribosomal DNA.

**Fig. 4.**
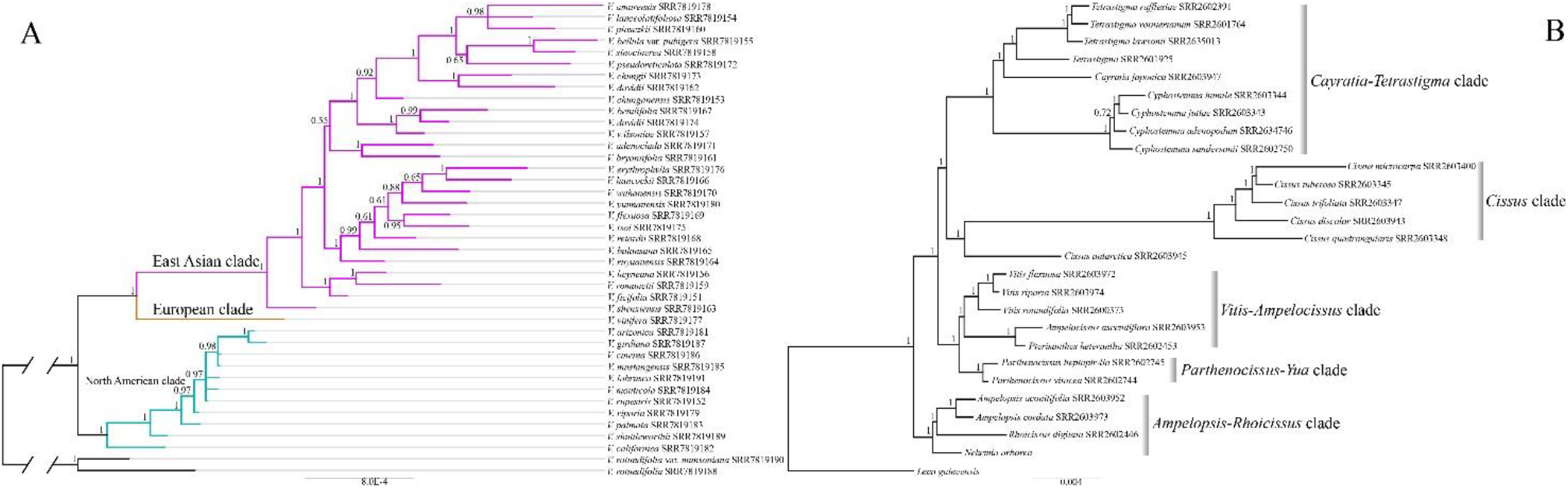
Bayesian trees inferred from 38 genes of (A) *Vitis* and (B) Vitaceae data. The number above the nodes indicate the branch support values measuring the support for the BI posterior probabilities (PP). Scale bars indicate substitutions per site.

As for the low-coverage Vitaceae data, the plastomes were well assembled using the successive reference approach except for some gaps in 10 samples, and this result was consistent with that in Zhang et al. (2015). Due to the ambiguous alignment in the intergenic regions of plastomes within the Vitaceae dataset, the plastid CDS regions were extracted for phylogenetic inference, and this data matrix can be accessed from Dryad Digital Repository Data S11: https://doi.org/10.5061/dryad.b2rbnzsd7 (Liu et al., 2021). All nodes have been strongly supported with BS 100 for ML and PP 1 for BS analysis (Fig. S8). In addition, the well-assembled nrDNA repeats from Vitaceae data (Table S2) resulted in a strongly supported phylogeny, and the five major clades recovered in the previous studies (Wen et al., 2013; Zhang et al., 2015) have been resolved in our nrDNA tree (Fig. 3B) except for the *Cissus* L. clade, in which the phylogenetic relationship between *C. antarctica* Vent. and the other four *Cissus* samples was not resolved (Fig. 3B). Our nrDNA tree also corroborated the sister relationship between the *Ampelopsis-Rhoicissus* clade and the other four clades, as well as between the *Parthenocissus* Planch. clade and *Vitis-Ampelocissus* clade. Furthermore, 38 target mitochondrial origin protein-coding genes were also successfully assembled, although with gaps for some samples (Dryad Digital Repository: Data S12, Liu et al., 2021). The mtDNA tree (Fig. 4B) resulted in a nearly similar topology to Zhang et al. (2015), but with greatly increased resolution than in Zhang et al. (2015)’s mtDNA tree (16 regions). For example, the monophyly of *Tetrastigma* (Miq.) Planch. is supported in our 38 mitochondrial gene data, and it was not supported due to insufficient characters in Zhang et al. (2015). All three phylogenies based on the plastome, mitochondrial genes, and nrDNA repeats resulted in nearly identical topologies except for the lower resolution of deep relationships among the five major clades in the nrDNA tree (Fig. 3B).

## 4 Discussion

### 4.1 Deep Genome Skimming (DGS) as an alternative to Hyb-Seq in Vitaceae

We recovered 618 SCNs from more than 50% samples of the 8× coverage data (Table 1 & Drayd Digital Repository Data S4), generating a large dataset for phylogenetic inference. However, the missing data in some sequences may prevent accurate inferences of phylogeny. With the decreased sequencing cost, especially on the Illumina or Novoseq platforms, it is practical to increase the sequencing depth of genome skimming. With a genome size of c. 500 MB (e.g., grapevine and apple), the 10× coverage is around 5 GB data, which cost c. $40 for each sample (NOVOgene, Beijing). Considering the balance between sufficient SCNs for phylogenetic inference and the cost, we suggest 10× coverage as the optimal sequencing depths for recovering SCNs. We herein propose a deep genome skimming (DGS) workflow (Fig. 5), that recovers SCNs, as well as the high-copy fraction of genomes: organelle genomes and nrDNA repeats.

**Fig. 5.**
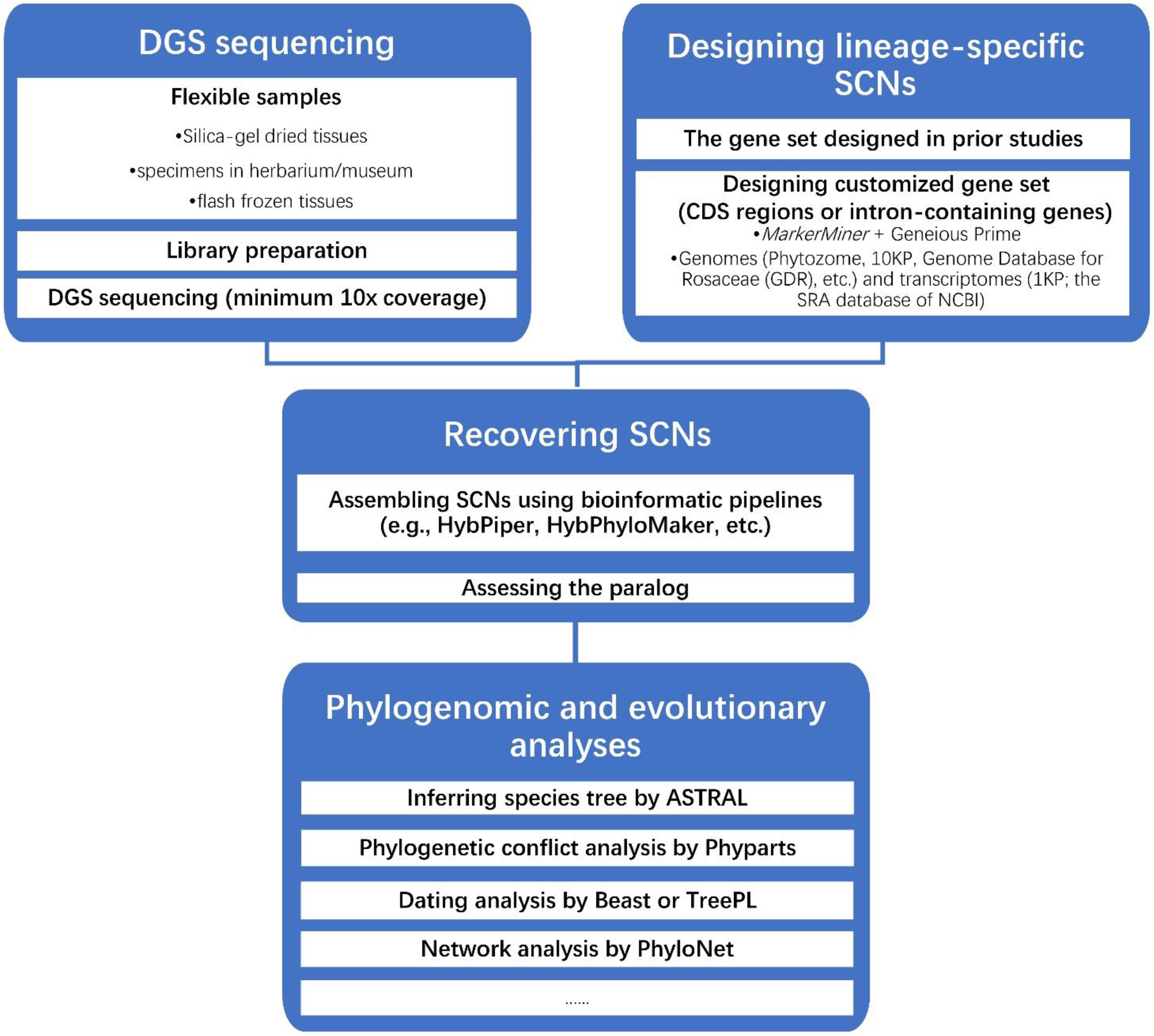
Illustrated workflow for designing and exploring single-copy nuclear genes from deep genome skimming data.

Contrasting to RADseq and RNAseq method, the DGS method can effectively utilize degraded DNA from herbarium specimens of rare, extinct, or ancient samples, which has been well tested in recent studies (Särkinen et al., 2012; Bakker et al., 2016; Saeidi et al., 2018; Liu et al., 2019, 2020a). Furthermore, given the ability to incorporate whole genomic information, DGS provides a possibility for assembling the intron-containing SCNs. The well-resolved gene trees estimated from the intron-containing SCNs will be useful for phylogenetic analyses, especially at the species level with relatively low sequence divergence. However, the intronic locus design for recovering SCNs requires a genomic reference in the study group (de Sousa et al., 2014; Weitemier et al., 2014) or a large number of contigs from high-coverage genome skimming data (Folk et al., 2015). DGS also provides the opportunity for plant systematists to capture different combinations of SCNs (e.g., with intron or without intron) to test the potential phylogenetic relationships among diverse lineages. Without the need for bait synthesis and target enrichment experiment, as well as the decreased sequencing cost, DGS is an economical approach to obtain large datasets of SCNs from non-model organisms and can serve as an alternative to Hyb-Seq.

Through simulation, the 10× coverage data in the *Vitis* case study has performed well in recovering lineage-specific SCNs (Fig. 2). The topology recovered from 10× coverage data (Fig. S2) was nearly similar to that from 20× coverage data (Fig. 2A), except for some nodes with lower support. We used the species tree estimated from the data matrix from the 20× coverage for the phylogenetic analysis. This case study based on 887 SCNs and whole plastomes provided sufficient informative sites for robustly constructing the phylogenetic relationships in *Vitis* (Fig. 2). North American *Vitis* subgenus *Vitis* (i.e., the North American clade in Figs. 2A, 4A) was supported to be monophyletic based on chloroplast and nuclear data in several previous studies (Jansen et al., 2006; Tröndle et al., 2010; Péros et al., 2011; Ren et al., 2011; Zecca et al., 2012; Aradhya et al., 2013; Miller et al., 2013; Liu et al., 2016; Ma et al., 2018; Wen et al., 2018a). This result was also supported in our plastid (Fig. 2B) and mtDNA (Fig. 4A) trees. However, the coalescent (ASTRAL species tree) and concatenated analyses (ML tree) of 887 SCNs both supported the paraphyly of the North American species of *Vitis* subgenus *Vitis* (Fig. 2A), consistent with trees from 27 single-copy nuclear markers (Wan et al., 2013), and the Hyb-Seq results (Nie et al., submitted). Of interest, the North American *Vitis* was monophyletic in the SNP phylogeny from Ma et al. (2018), although these two different data matrices (SNPs and 887 SCNs) have been generated from the same raw data. Furthermore, *Vitis labrusca* was placed sister to the European species, *V. vinifera* in our SCN topologies (Fig. 2A), but it grouped with the North American species in the plastid (Fig. 2B) and mtDNA (Fig. 4A) trees. Because the tissue sample of *V. labrusca* was obtained as a cultivar in Henan, China in Ma et al. (2018), our results suggest that the sample likely represents the hybrid between *V. labrusca* and *V. vinifera*. Although the coalescent and concatenated analyses of 887 SCNs resulted in highly supported values for each node, the *phyparts* analyses showed that most of the pies (Fig. 2A) were filled with gray areas, indicating that there was limited support from only a few resolved nuclear genes. This result may be explained by the insufficient informative sites to resolve the species relationships within *Vitis* for each nuclear locus. Incorporating intron regions for each gene in the future may help provide more informative characters to overcome this problem.

An important step in successfully utilizing the DGS method is to design the potential SCNs for the studied lineages (Fig. 5). It has become straightforward to design single/low-copy nuclear gene marker sets using available transcriptomes or whole genomes, for capturing SCNs *in silico* as we have advocated here for the DGS method. According to the database of plaBi-PD (https://plabipd.de/plant_genomes_pa.ep; data of accession: Jan. 22, 2020), a total of 498 Angiosperm genomes have been published covering 42 orders and 107 families. In addition, there are numerous transcriptome sequences available (e.g., the 1KP Project; https://sites.google.com/a/ualberta.ca/onekp/; the SRA database of NCBI). These are great resources for the selection and development of nuclear gene markers to address specific questions, and these resources are accumulating at a rapid and increasing rate every year. For example, by the end of 2022, over 10,000 plant genomes may be sequenced, representing all major clades of plants and eukaryotic microbes via the 10,000 Plant Genomes Project (10KP; https://db.cngb.org/10kp) (Cheng et al., 2018). Pipelines or programs for target loci selection have also been developed (Weitemier et al., 2014; Yang & Smith, 2014; Chamala et al., 2015; Smith et al., 2020). Among others, MarkerMiner (Chamala et al., 2015) is a good starting tool to explore gene selection. It compares user-provided transcriptomic or genomic data against reference databases of known single-copy nuclear genes identified based on a survey of duplication-resistant genes in 17 angiosperm genomes by De Smet et al. (2013). It is easy to implement and also saves time by automating the selection process. However, since only 17 angiosperm genomes are compared by MarkerMiner, it will be useful to further filter the MarkerMiner-selected genes by comparing them in Geneious Prime (Kearse et al., 2012) against other available genomes or transcriptomes closely related to the groups of interest. Generally, a stringent selection criterion will facilitate the downstream gene assembly and phylogenetic analyses using Geneious Prime (Kearse et al., 2012) and/or the script *GoldFinder* developed by Vargas et al. (2019). Although universal single-copy nuclear gene sets have been developed, such as the Angiosperm-353 generic baits set (Johnson et al., 2019), several studies have compared the success of taxon-specific and universal SCNs and have found that taxon-specific single-copy nuclear loci dataset yield a higher number of phylogenetically informative loci (Kadlec et al., 2017; Chau et al., 2018; Jantzen et al., 2020; Straub et al., 2020). When feasible, we recommend a combination of lineage-specific and universal SCNs sets to yield the largest pool of nuclear loci appropriate for phylogenomic studies (also see Jantzen et al., 2020).

### 4.2 On the utility of the plastid genomes, nrDNA, and mtDNA

The nrDNA sequences of the family Vitaceae assembled here provide an example of generating the entire rDNA repeats, even from lower coverage genome skimming data (less than 6× coverage data) in Vitaceae. The five major clades (Fig. 3B) supported by transcriptomic (Wen et al., 2013) and Hyb-Seq (Ma et al., 2021) data have also been recovered from the nrDNA data, indicating the great value of nrDNA in phylogenetic inference. Nevertheless, for nrDNA studies in the past, e.g., the intragenomic polymorphisms among nrDNA repeats likely arising from incomplete concerted evolution have limited their wide utilization in plant systematics and may affect the precise estimates of branch length or divergence times (Weitemier et al., 2015; Fonseca & Lohmann, 2020). Nevertheless, nrDNA sequences will continue to be an important resource for inferring plant phylogeny with easy access to the entire nrDNA repeats from genome skimming, but the extent of intragenomic polymorphisms needs to be evaluated rigorously.

Mitochondrial DNA has not been broadly utilized for phylogenetic analyses in plants relative to the nuclear and plastid genomes, because of its low nucleotide substitution rates and potentially evolutionary history shared with the plastome (Wolfe et al., 1987; Sloan et al., 2009; Fonseca & Lohmann, 2020). However, our case studies of mtDNA either at the genus level (*Vitis*) or at the family level (Vitaceae) have provided additional information/insights into the phylogenetic relationships among lineages. The mtDNA tree based on 38 genes resulted in a well-supported backbone of Vitaceae, in which the monophyly of *Tetrastigma* and the sister relationship between *Cayratia* Juss.+*Tetrastigma* and *Cyphostemma* (Planch.) Alston. (Fig. 4B) were supported with the incorporation of more mitochondrial origin genes (38 genes). The utility of mtDNA at the family level (Fig. 4B) was also corroborated in the recent case study of Rubiaceae (Rydin et al., 2017). Additionally, several studies have shown the successful utilization of mtDNA in deep-level phylogenetics, such as among 280 genera of angiosperms (Adams et al., 2002) and at the ordinal level of mosses (Beckert et al., 2001). Additionally, our mtDNA tree at the genus-level case study of *Vitis* resulted in three strongly supported clades and provided some insights into the phylogenetic relationships within *Vitis* (Fig. 4A). However, mtDNA in general has not been a highly informative phylogenetic marker at the genus level (Galtier et al., 2009; Spooner et al., 2020a, 2020b)due to the low sequence substitution rate.

## 5 Conclusions

Genome skimming with low-coverage sequencing depth (less than 5× coverage) has proven to be successful and economical for assembling plastome, genic portion of mtDNA, and entire nrDNA repeats (Straub et al., 2012; Fonseca & Lohmann, 2020). The drastic decrease in sequencing cost in recent years has provided a good opportunity for plant systematists to obtain more data at an affordable cost. Our simulations and empirical results demonstrate that 10× coverage data enables capturing of sufficiently customized SCNs datasets for downstream phylogenetic and evolutionary analysis in the case study of *Vitis*. Our comparative results showed the efficacy of assembling the entire nrDNA repeats sequences from genome skimming data and its significance in phylogenetic inference. The well-assembled mtDNA also showed great promise in reconstructing phylogenetic relationships at the higher taxonomic levels, particularly at the family level, while mtDNA can also provide some meaningful insights into some species relationships. The increased sequencing depth of genome skimming (i.e., DGS) will facilitate the elucidation of phylogenetic relationships in nonmodel organisms. DGS can economically capture large datasets of SCNs *in silico* without the need to synthesize baits and to use complicated enrichment experiments.

## Acknowledgments

All computational analyses were conducted on the Smithsonian Institution High Performance Computing Cluster (SI/HPC, “Hydra”). This research is supported by the National Natural Science Foundation of China (Grant No.: 32000163 & 31620103902).

---

**Table S1.**
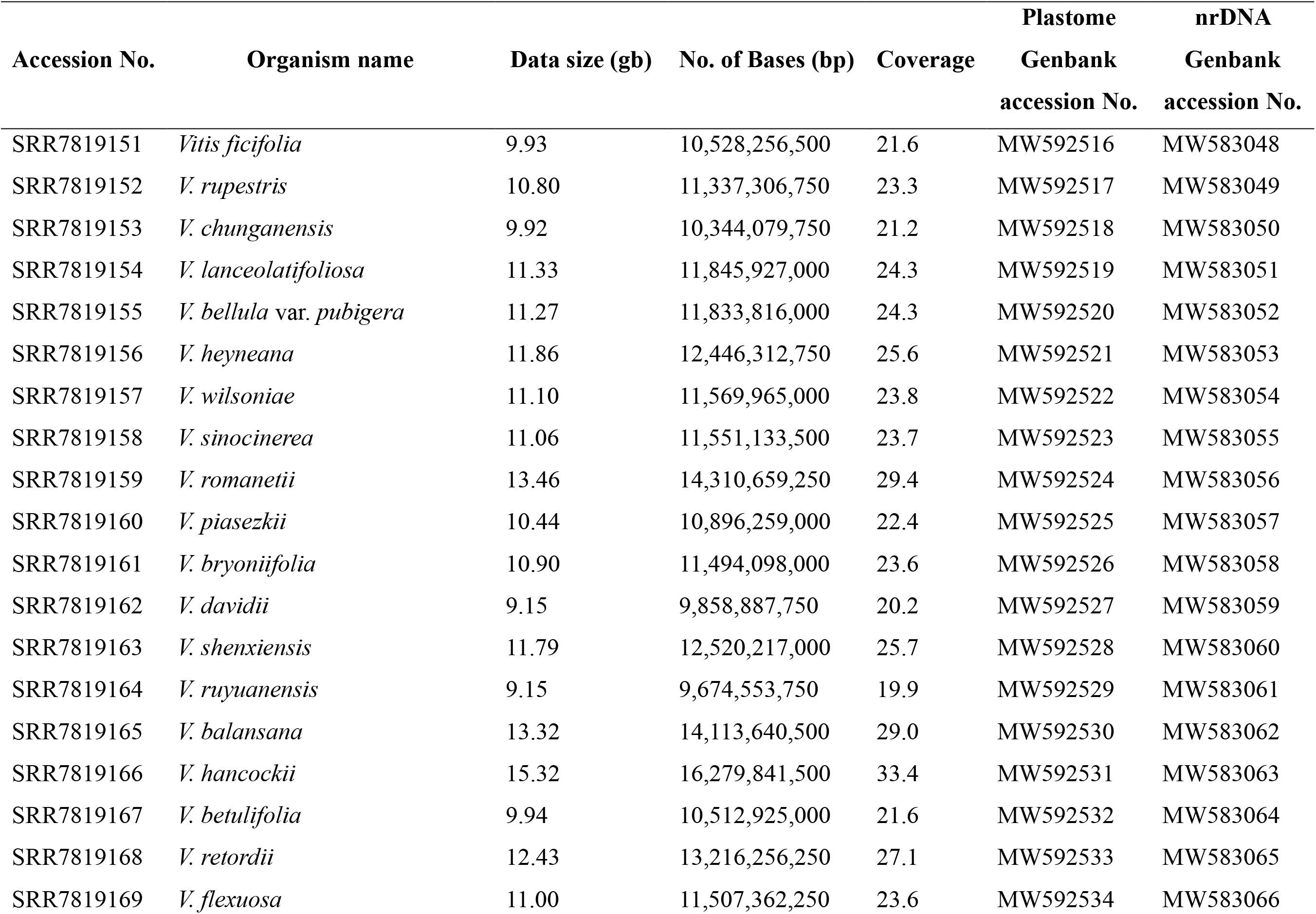
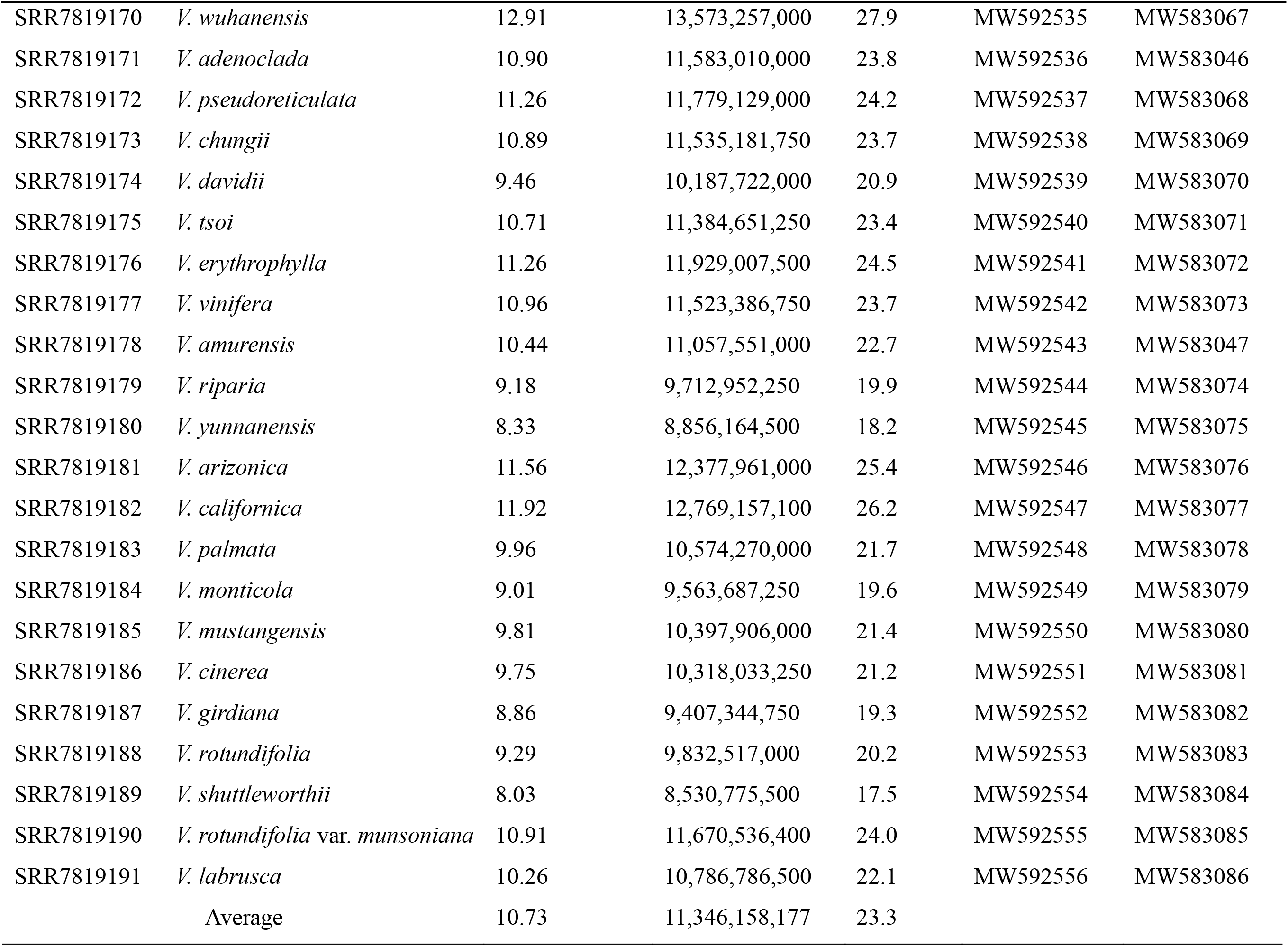
Sampling and sequencing information for the genus-level case (*Vitis* data)

| Accession No. | Organism name | Data size (gb) | No. of Bases (bp) | Coverage | Plastome | nrDNA |
| --- | --- | --- | --- | --- | --- | --- |
|  |  |  |  |  | Genbank<br>accession No. | Genbank<br>accession No. |
| SRR7819151 | *Vitis ficifolia* | 9.93 | 10,528,256,500 | 21.6 | MW592516 | MW583048 |
| SRR7819152 | *V. rupestris* | 10.80 | 11,337,306,750 | 23.3 | MW592517 | MW583049 |
| SRR7819153 | *V. chunganensis* | 9.92 | 10,344,079,750 | 21.2 | MW592518 | MW583050 |
| SRR7819154 | *V. lanceolatifolia* | 11.33 | 11,845,927,000 | 24.3 | MW592519 | MW583051 |
| SRR7819155 | *V. bellula* var. *pubigera* | 11.27 | 11,833,816,000 | 24.3 | MW592520 | MW583052 |
| SRR7819156 | *V. heyneana* | 11.86 | 12,446,312,750 | 25.6 | MW592521 | MW583053 |
| SRR7819157 | *V. wilsoniae* | 11.10 | 11,569,965,000 | 23.8 | MW592522 | MW583054 |
| SRR7819158 | *V. sinocinerea* | 11.06 | 11,551,133,500 | 23.7 | MW592523 | MW583055 |
| SRR7819159 | *V. romanetii* | 13.46 | 14,310,659,250 | 29.4 | MW592524 | MW583056 |
| SRR7819160 | *V. piasezkii* | 10.44 | 10,896,259,000 | 22.4 | MW592525 | MW583057 |
| SRR7819161 | *V. bryoniifolia* | 10.90 | 11,494,098,000 | 23.6 | MW592526 | MW583058 |
| SRR7819162 | *V. davidii* | 9.15 | 9,858,887,750 | 20.2 | MW592527 | MW583059 |
| SRR7819163 | *V. shenxiensis* | 11.79 | 12,520,217,000 | 25.7 | MW592528 | MW583060 |
| SRR7819164 | *V. ruyuanensis* | 9.15 | 9,674,553,750 | 19.9 | MW592529 | MW583061 |
| SRR7819165 | *V. balansana* | 13.32 | 14,113,640,500 | 29.0 | MW592530 | MW583062 |
| SRR7819166 | *V. hancockii* | 15.32 | 16,279,841,500 | 33.4 | MW592531 | MW583063 |
| SRR7819167 | *V. betulifolia* | 9.94 | 10,512,925,000 | 21.6 | MW592532 | MW583064 |
| SRR7819168 | *V. retordii* | 12.43 | 13,216,256,250 | 27.1 | MW592533 | MW583065 |
| SRR7819169 | *V. flexuosa* | 11.00 | 11,507,362,250 | 23.6 | MW592534 | MW583066 |
| SRR7819170 | *V. wuhanensis* | 12.91 | 13,573,257,000 | 27.9 | MW592535 | MW583067 |
| SRR7819171 | *V. adenoclada* | 10.90 | 11,583,010,000 | 23.8 | MW592536 | MW583046 |
| SRR7819172 | *V. pseudoreticulata* | 11.26 | 11,779,129,000 | 24.2 | MW592537 | MW583068 |
| SRR7819173 | *V. chungii* | 10.89 | 11,535,181,750 | 23.7 | MW592538 | MW583069 |
| SRR7819174 | *V. davidii* | 9.46 | 10,187,722,000 | 20.9 | MW592539 | MW583070 |
| SRR7819175 | *V. tsoi* | 10.71 | 11,384,651,250 | 23.4 | MW592540 | MW583071 |
| SRR7819176 | *V. erythrophylla* | 11.26 | 11,929,007,500 | 24.5 | MW592541 | MW583072 |
| SRR7819177 | *V. vinifera* | 10.96 | 11,523,386,750 | 23.7 | MW592542 | MW583073 |
| SRR7819178 | *V. amurensis* | 10.44 | 11,057,551,000 | 22.7 | MW592543 | MW583047 |
| SRR7819179 | *V. riparia* | 9.18 | 9,712,952,250 | 19.9 | MW592544 | MW583074 |
| SRR7819180 | *V. yunnanensis* | 8.33 | 8,856,164,500 | 18.2 | MW592545 | MW583075 |
| SRR7819181 | *V. arizonica* | 11.56 | 12,377,961,000 | 25.4 | MW592546 | MW583076 |
| SRR7819182 | *V. californica* | 11.92 | 12,769,157,100 | 26.2 | MW592547 | MW583077 |
| SRR7819183 | *V. palmata* | 9.96 | 10,574,270,000 | 21.7 | MW592548 | MW583078 |
| SRR7819184 | *V. monticola* | 9.01 | 9,563,687,250 | 19.6 | MW592549 | MW583079 |
| SRR7819185 | *V. mustangensis* | 9.81 | 10,397,906,000 | 21.4 | MW592550 | MW583080 |
| SRR7819186 | *V. cinerea* | 9.75 | 10,318,033,250 | 21.2 | MW592551 | MW583081 |
| SRR7819187 | *V. girdiana* | 8.86 | 9,407,344,750 | 19.3 | MW592552 | MW583082 |
| SRR7819188 | *V. rotundifolia* | 9.29 | 9,832,517,000 | 20.2 | MW592553 | MW583083 |
| SRR7819189 | *V. shuttleworthii* | 8.03 | 8,530,775,500 | 17.5 | MW592554 | MW583084 |
| SRR7819190 | *V. rotundifolia* var. *munsoniana* | 10.91 | 11,670,536,400 | 24.0 | MW592555 | MW583085 |
| SRR7819191 | *V. labrusca* | 10.26 | 10,786,786,500 | 22.1 | MW592556 | MW583086 |
|  | Average | 10.73 | 11,346,158,177 | 23.3 |  |  |

**Table S2.**
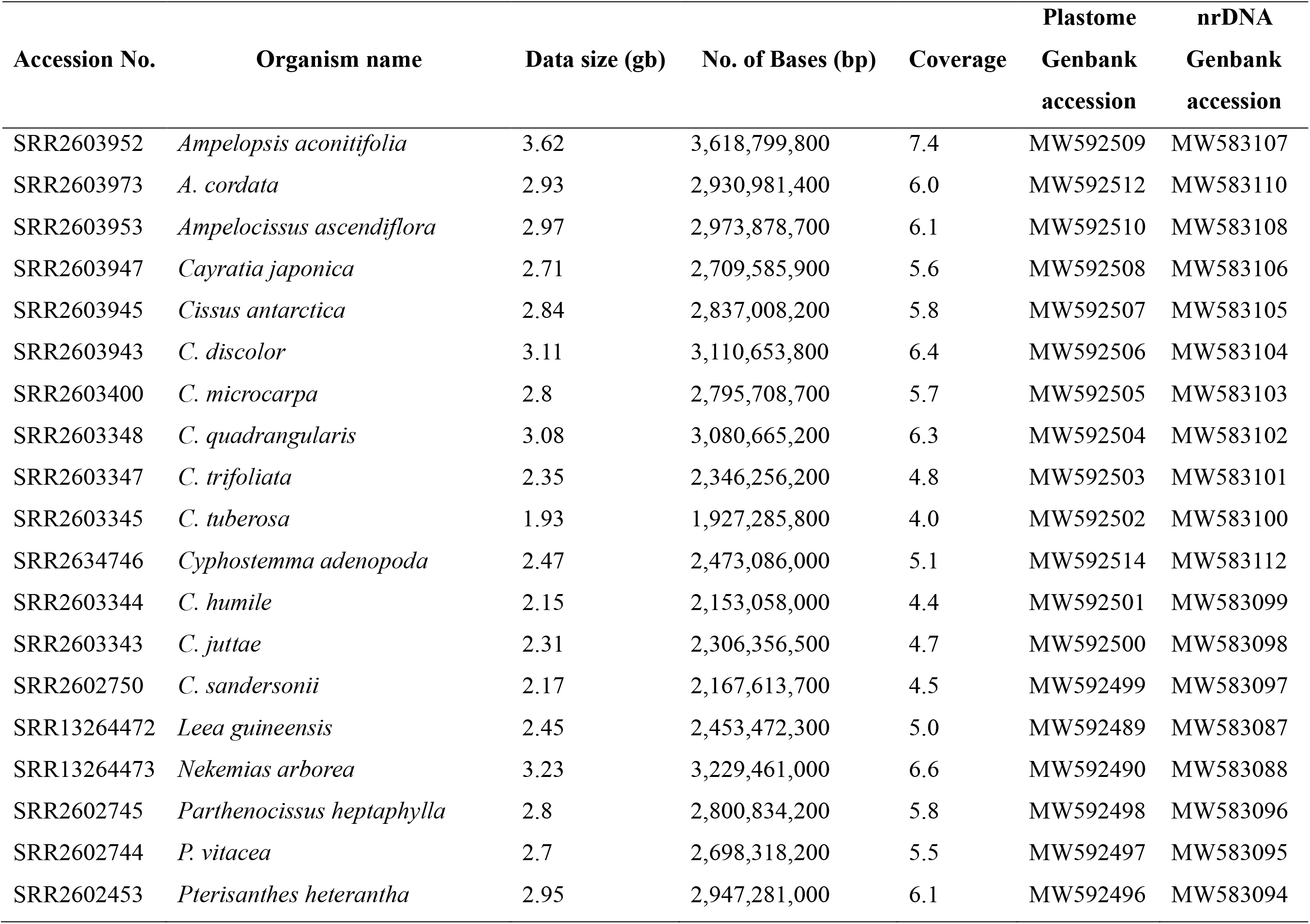
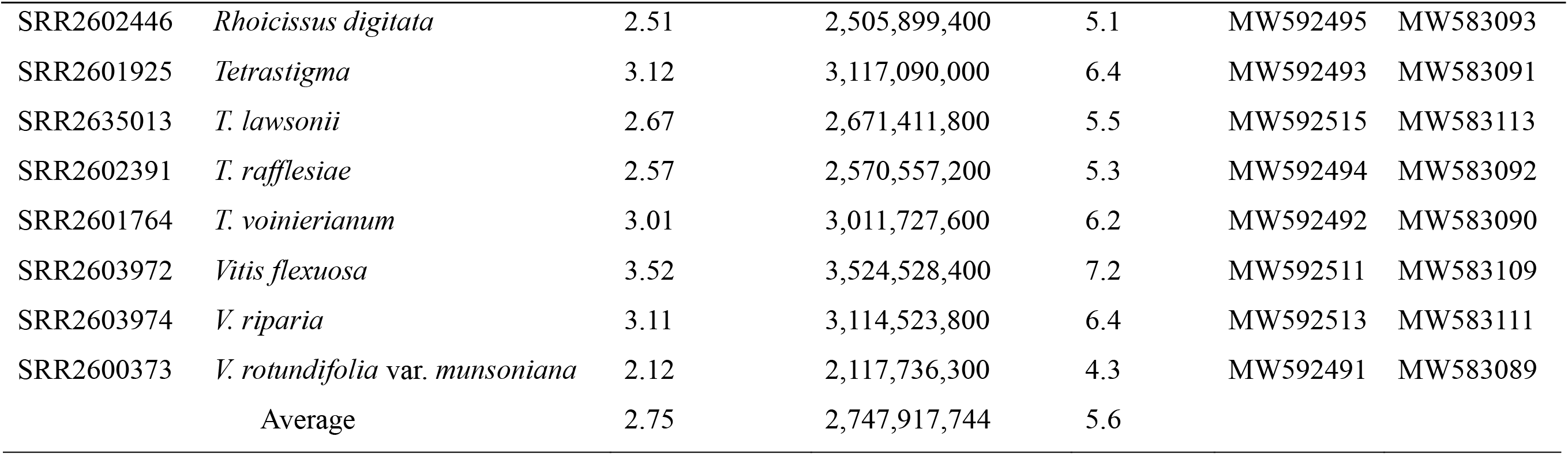
Sampling and sequencing information of the family-level case (Vitaceae data)

| Accession No. | Organism name | Data size (gb) | No. of Bases (bp) | Coverage | Plastome | nrDNA |
| --- | --- | --- | --- | --- | --- | --- |
|  |  |  |  |  | Genbank<br>accession | Genbank<br>accession |
| SRR2603952 | *Ampelopsis aconitifolia* | 3.62 | 3,618,799,800 | 7.4 | MW592509 | MW583107 |
| SRR2603973 | *A. cordata* | 2.93 | 2,930,981,400 | 6.0 | MW592512 | MW583110 |
| SRR2603953 | *Ampelocissus ascendiflora* | 2.97 | 2,973,878,700 | 6.1 | MW592510 | MW583108 |
| SRR2603947 | *Cayratia japonica* | 2.71 | 2,709,585,900 | 5.6 | MW592508 | MW583106 |
| SRR2603945 | *Cissus antarctica* | 2.84 | 2,837,008,200 | 5.8 | MW592507 | MW583105 |
| SRR2603943 | *C. discolor* | 3.11 | 3,110,653,800 | 6.4 | MW592506 | MW583104 |
| SRR2603400 | *C. microcarpa* | 2.8 | 2,795,708,700 | 5.7 | MW592505 | MW583103 |
| SRR2603348 | *C. quadrangularis* | 3.08 | 3,080,665,200 | 6.3 | MW592504 | MW583102 |
| SRR2603347 | *C. trifoliata* | 2.35 | 2,346,256,200 | 4.8 | MW592503 | MW583101 |
| SRR2603345 | *C. tuberosa* | 1.93 | 1,927,285,800 | 4.0 | MW592502 | MW583100 |
| SRR2634746 | *Cyphostemma adenopoda* | 2.47 | 2,473,086,000 | 5.1 | MW592514 | MW583112 |
| SRR2603344 | *C. humile* | 2.15 | 2,153,058,000 | 4.4 | MW592501 | MW583099 |
| SRR2603343 | *C. juttae* | 2.31 | 2,306,356,500 | 4.7 | MW592500 | MW583098 |
| SRR2602750 | *C. sandersonii* | 2.17 | 2,167,613,700 | 4.5 | MW592499 | MW583097 |
| SRR13264472 | *Leea guineensis* | 2.45 | 2,453,472,300 | 5.0 | MW592489 | MW583087 |
| SRR13264473 | *Nekemias arborea* | 3.23 | 3,229,461,000 | 6.6 | MW592490 | MW583088 |
| SRR2602745 | *Parthenocissus heptaphylla* | 2.8 | 2,800,834,200 | 5.8 | MW592498 | MW583096 |
| SRR2602744 | *P. vitacea* | 2.7 | 2,698,318,200 | 5.5 | MW592497 | MW583095 |
| SRR2602453 | *Pterisanthes heterantha* | 2.95 | 2,947,281,000 | 6.1 | MW592496 | MW583094 |
| SRR2602446 | *Rhoicissus digitata* | 2.51 | 2,505,899,400 | 5.1 | MW592495 | MW583093 |
| SRR2601925 | *Tetrastigma* | 3.12 | 3,117,090,000 | 6.4 | MW592493 | MW583091 |
| SRR2635013 | *T. lawsonii* | 2.67 | 2,671,411,800 | 5.5 | MW592515 | MW583113 |
| SRR2602391 | *T. rafflesiae* | 2.57 | 2,570,557,200 | 5.3 | MW592494 | MW583092 |
| SRR2601764 | *T. voinierianum* | 3.01 | 3,011,727,600 | 6.2 | MW592492 | MW583090 |
| SRR2603972 | *Vitis flexuosa* | 3.52 | 3,524,528,400 | 7.2 | MW592511 | MW583109 |
| SRR2603974 | *V. riparia* | 3.11 | 3,114,523,800 | 6.4 | MW592513 | MW583111 |
| SRR2600373 | *V. rotundifolia* var. *munsoniana* | 2.12 | 2,117,736,300 | 4.3 | MW592491 | MW583089 |
|  | Average | 2.75 | 2,747,917,744 | 5.6 |  |  |

## Supplementary Material

The following supplementary material is available online for this article at…

**Fig. S1.** ASTRAL species tree inferred from 31 SCNs assembled from the 6× coverage genome skimming data of *Vitis* in silico. The number above the nodes indicate the branch support values measuring the support for a local posterior possibility.

**Fig. S2.** ASTRAL species tree inferred from 618 SCNs assembled from the 8× coverage genome skimming data of *Vitis* in silico. The number above the nodes indicate the branch support values measuring the support for a local posterior possibility.

**Fig. S3.** ASTRAL species tree inferred from 876 SCNs assembled from the 10× coverage genome skimming data of *Vitis* in silico. The number above the nodes indicate the branch support values measuring the support for a local posterior possibility.

**Fig. S4.** ASTRAL species tree inferred from 885 SCNs assembled from the 12× coverage genome skimming data of *Vitis* in silico. The number above the nodes indicate the branch support values measuring the support for a local posterior possibility.

**Fig. S5.** ASTRAL species tree inferred from 887 SCNs assembled from the 14× coverage genome skimming data of *Vitis* in silico. The number above the nodes indicate the branch support values measuring the support for a local posterior possibility.

**Fig. S6.** ASTRAL species tree inferred from 887 SCNs assembled from the 16× coverage genome skimming data of *Vitis* in silico. The number above the nodes indicate the branch support values measuring the support for a local posterior possibility.

**Fig. S7** ASTRAL species tree inferred from 887 SCNs assembled from the 18× coverage genome skimming data of *Vitis* in silico. The number above the nodes indicate the branch support values measuring the support for a local posterior possibility.

**Fig. S8** Bayesian trees inferred from 80 plastid coding sequences (CDS) of Vitaceae data. The number above the nodes indicate the branch support values measuring the support for the BI posterior probabilities (PP), and all nodes have PP values of 1 unless noted otherwise. Scale bars indicate substitutions per site.

