## Supplementary figures and images for "Capturing single-copy nuclear genes, organellar genomes, and nuclear ribosomal DNA from deep genome skimming data for plant phylogenetics: A case study in Vitaceae"

### Fig. S1

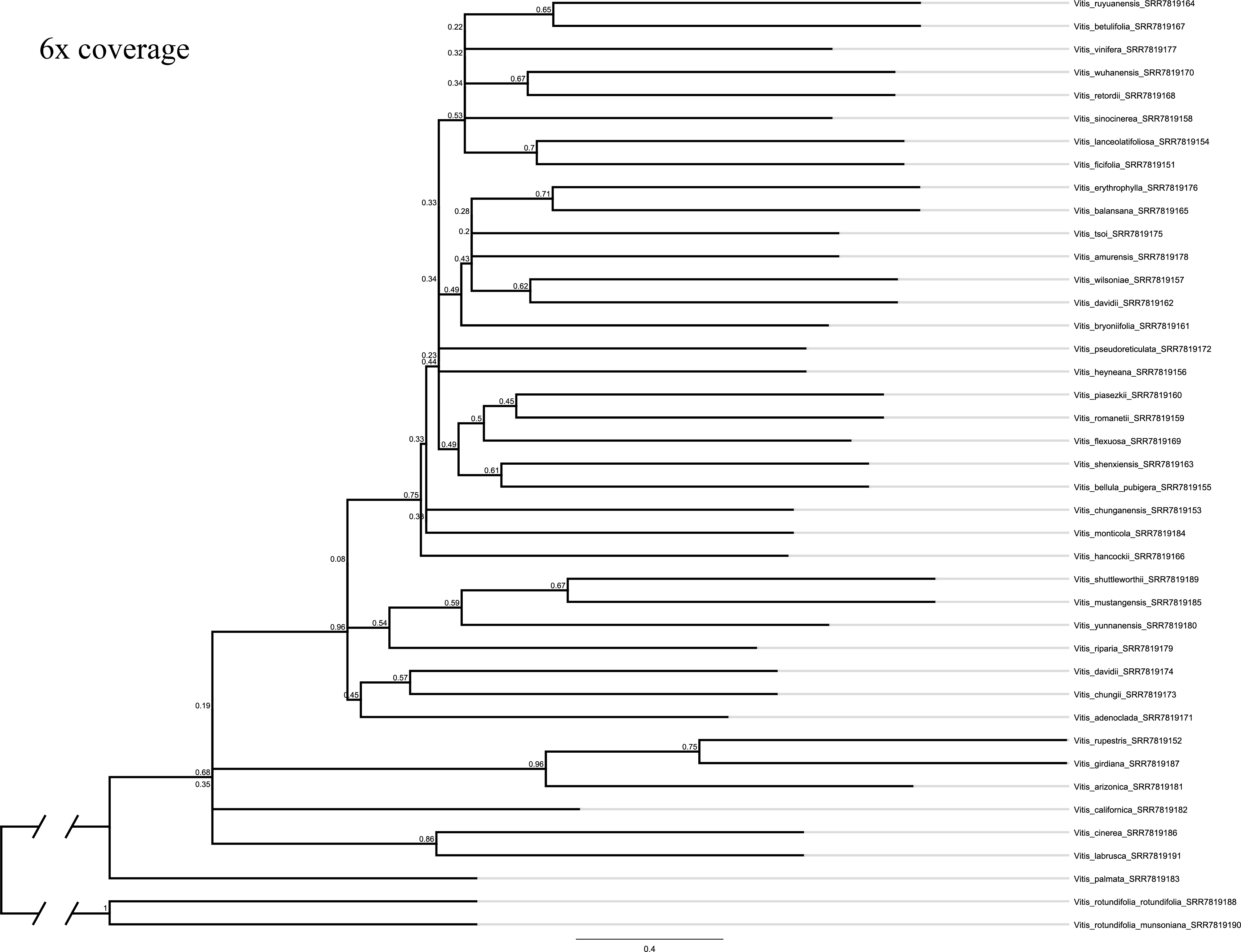

### Fig. S2

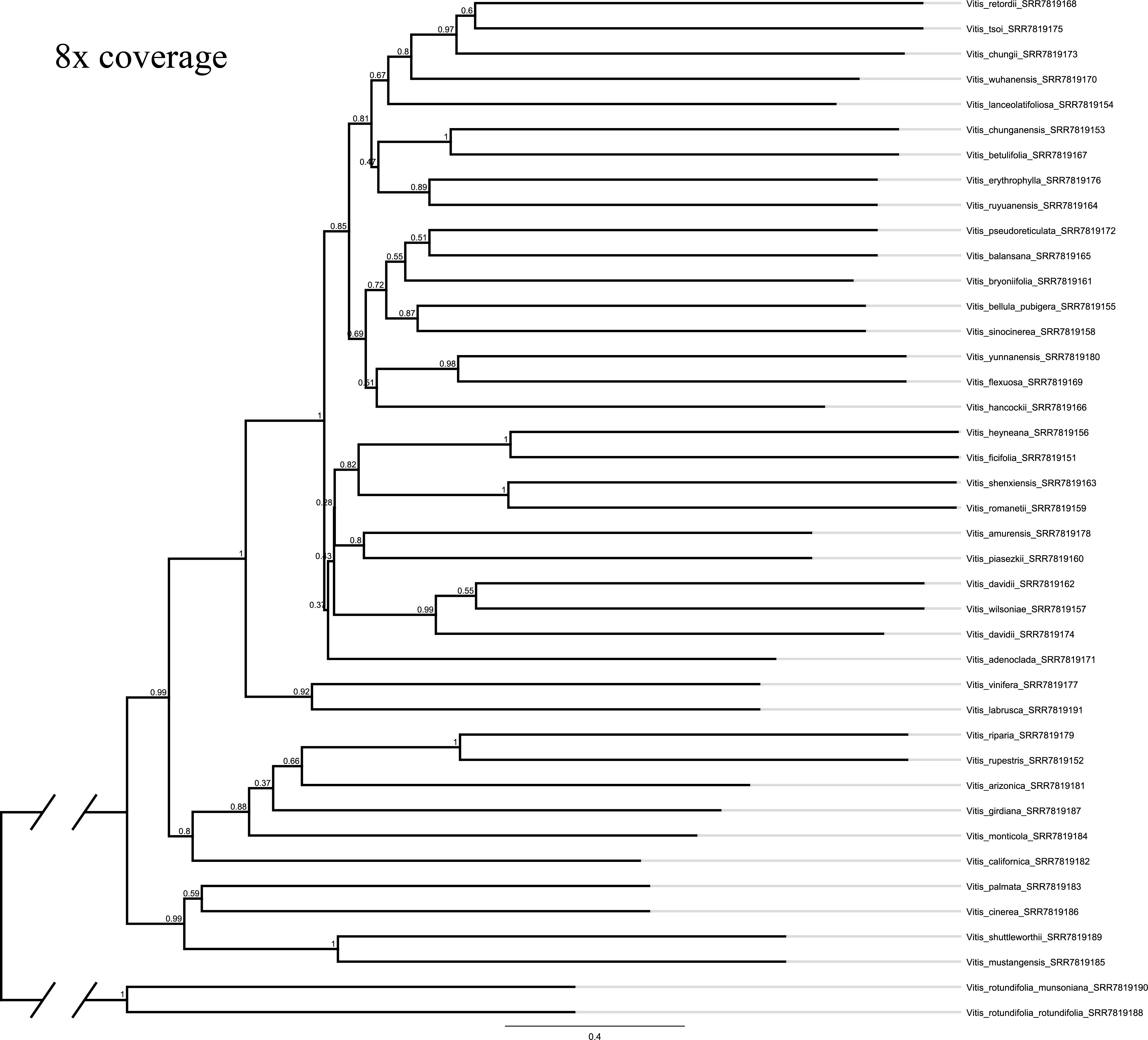

### Fig. S3

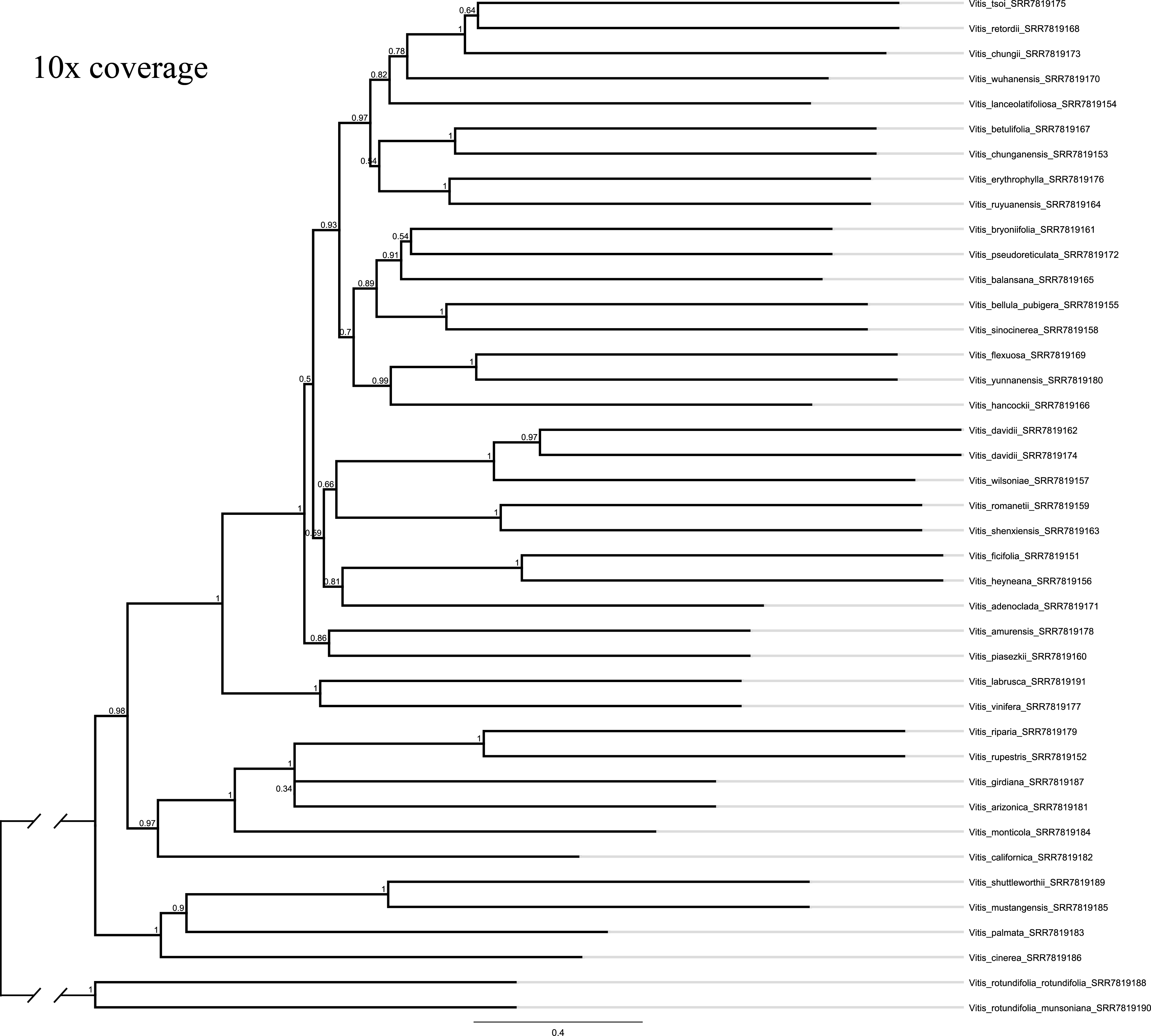

### Fig. S4

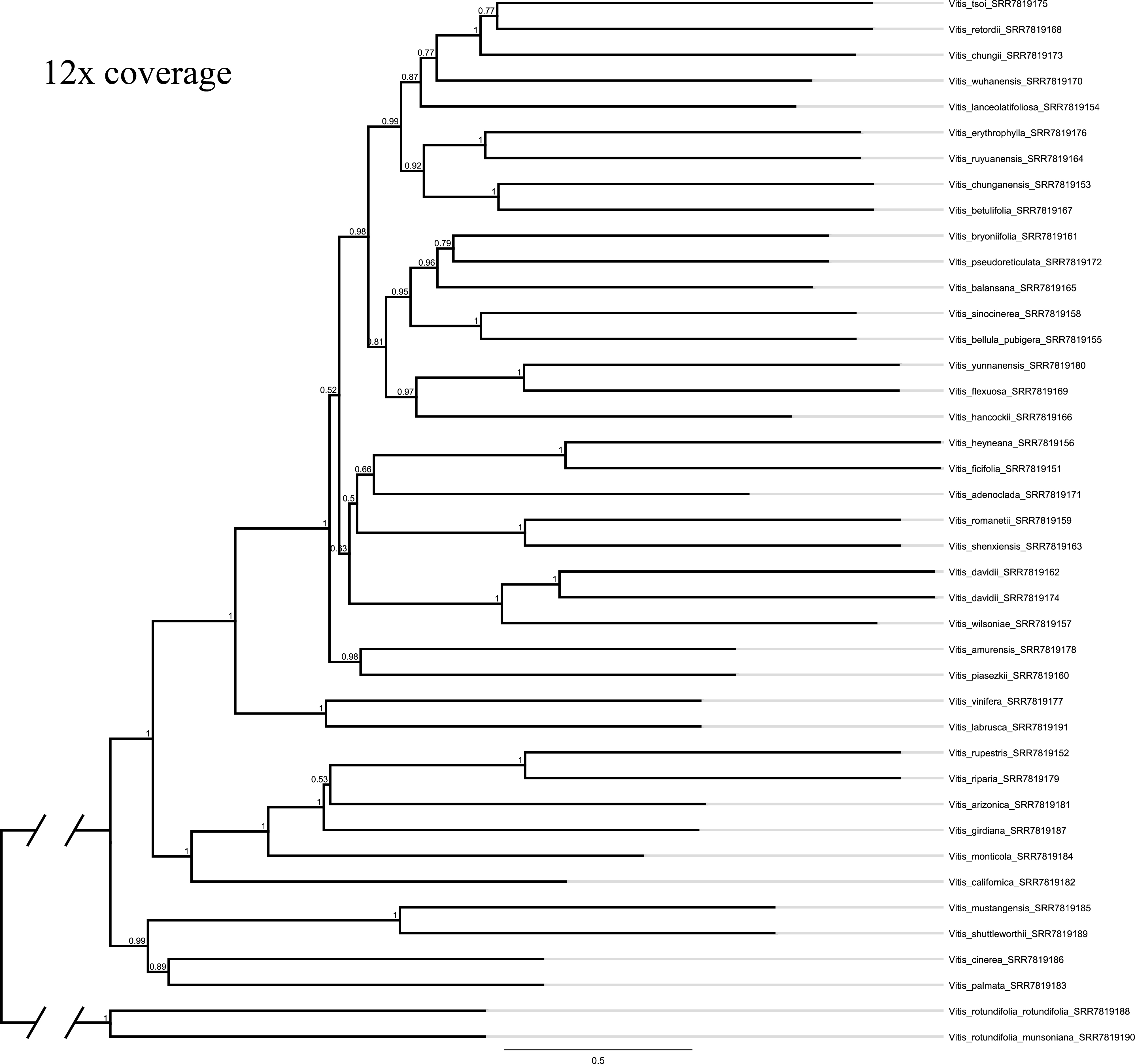

### Fig. S5

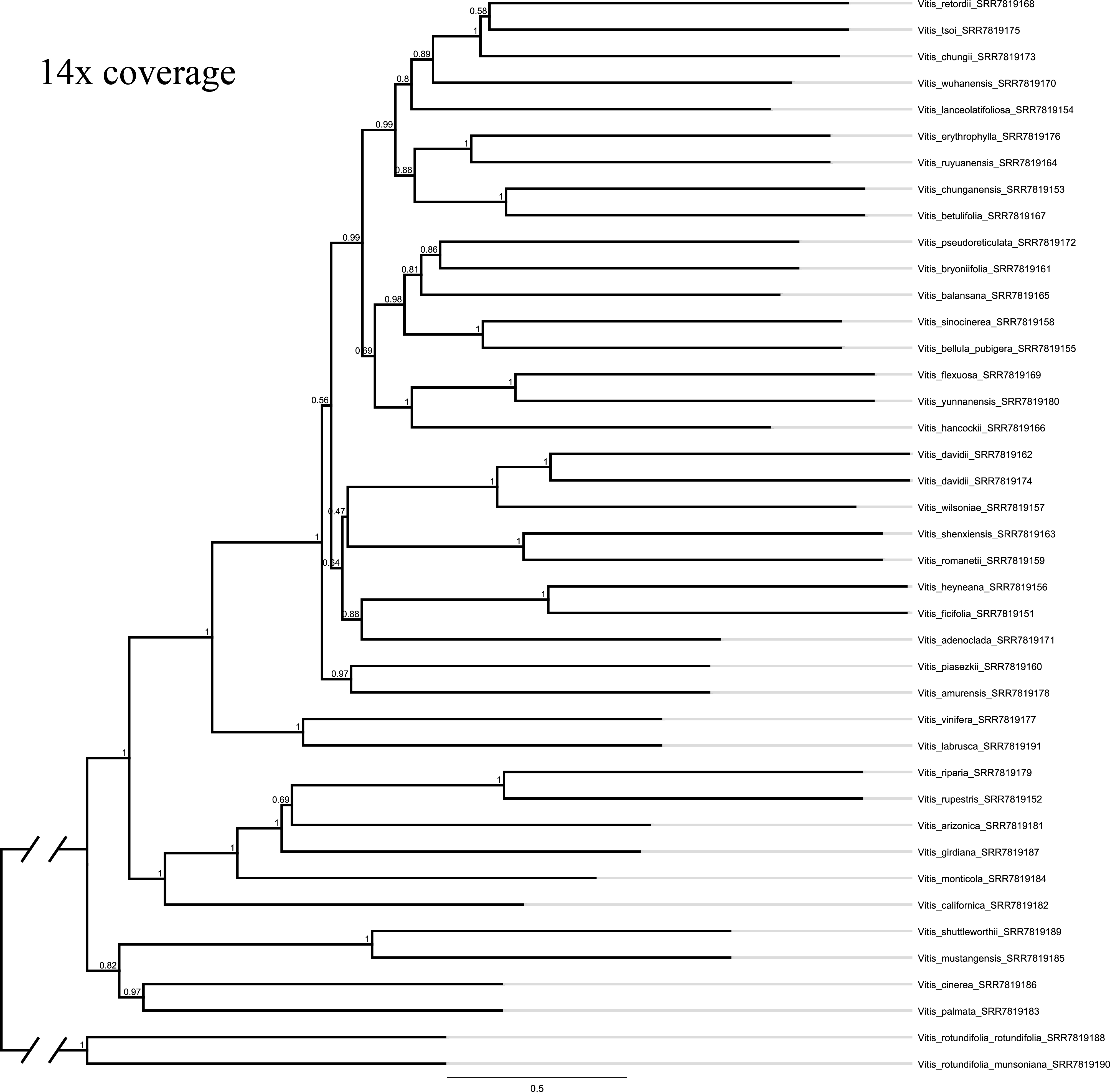

### Fig. S6

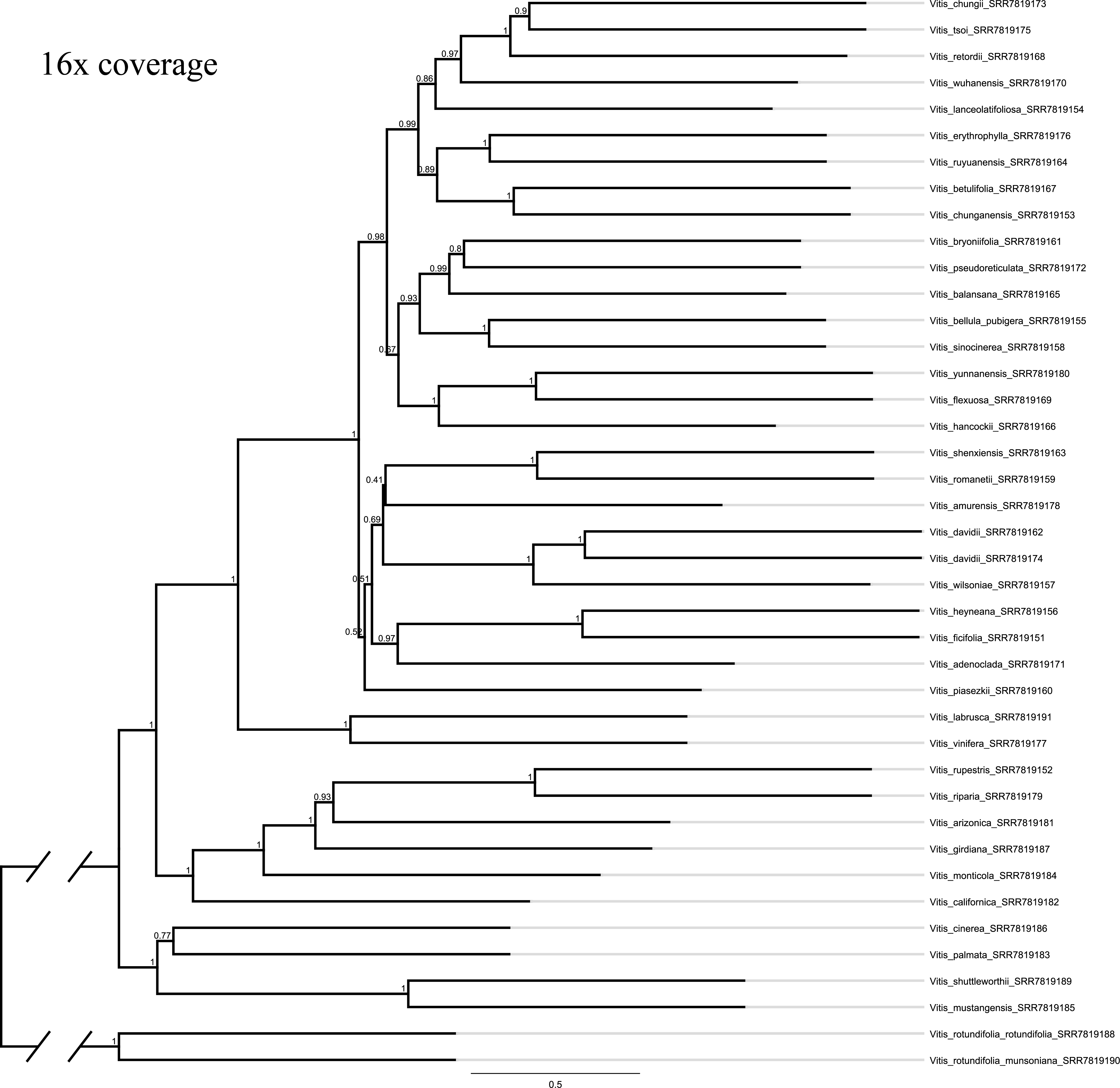

### Fig. S7

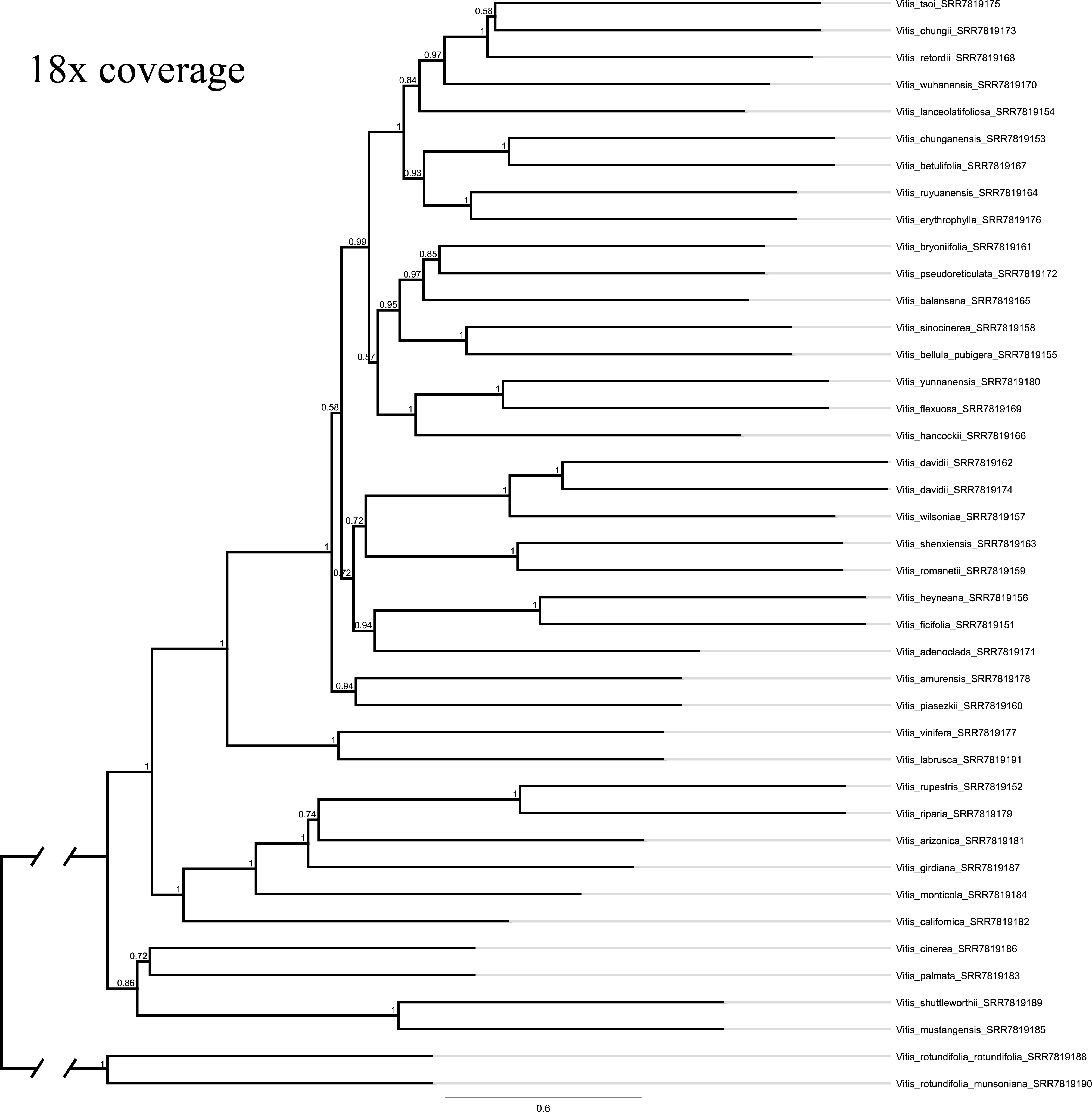

### Fig. S8

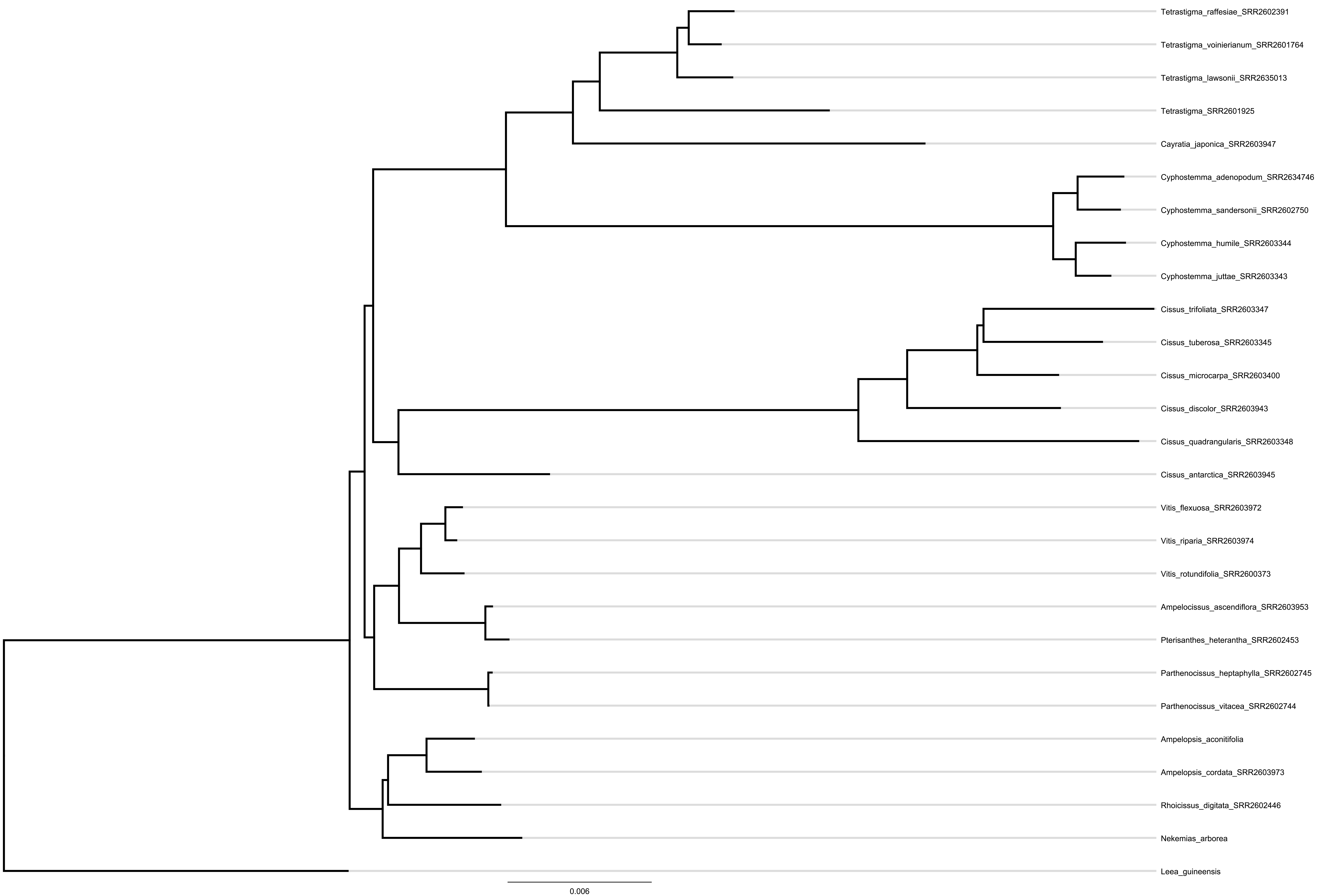
